# Hurdles to Horizontal Gene Transfer: Synonymous variation determines antibiotic resistance phenotype across species

**DOI:** 10.1101/2024.01.28.576264

**Authors:** Michael Finnegan, Caroline J Rose, Jeanne Hamet, Benjamin Prat, Stéphanie Bedhomme

## Abstract

Evidence that synonymous mutations and synonymous gene variants have fitness effects have accumulated recently. Since horizontal gene transfer represents a change in the genome of expression of the transferred gene, we hypothesized that the codon usage preferences of a horizontally transferred gene could determine the conferred fitness advantage or disadvantage, condition the immediate success of the transfer and in the longer term orient transfers.

To test this hypothesis, we characterized resistance levels of synonymous variants of a gentamicin resistance gene, inserted into a broad-host range plasmid and transformed into three different bacterial species *Escherichia coli, Acinetobacter baylyi and Pseudomonas aeruginosa.* We revealed a strong species effect, explained in part by differences in plasmid copy number between host species. Importantly, the relative levels of resistance conferred by each synonymous variant were not conserved across species, indicating that these phenotypic effects are due to differing compatibility between the transferred variants and the receiver bacterial genomes. This species-variant interaction confirms that the codon composition of a gene can be a determinant of post-horizontal gene transfer success. However, the similarity in codon usage between the synonymous variants and the host genome only explained the phenotypic differences between variants in one species, *P. aeruginosa*. Further investigations of the effects of local codon usage, translation bottlenecks and internal Shine-Dalgarno sequences did not reveal common universal mechanisms across our three bacterial species and point to multiple paths leading from the synonymous sequence to phenotype and a species-specific sensitivity to these different paths.

## Introduction

Horizontal gene transfer (HGT) plays a major role in bacterial evolution. It provides genetic variation, allowing bacterial populations to adapt to changing environments (Arnold et al. 2022). HGT rates vary amongst bacterial species and in some cases these HGT events occur at a greater rate than mutation, underlining their importance as drivers of evolution (Didelot et al. 2013; Frazão et al. 2019). Unlike most eukaryotes, which seldom undertake HGT, bacteria can potentially source fully fledged genes and other genetic materials from any other bacterium (Brito 2021). However, the horizontal spread of genetic material is not equivalent to simple diffusion and preferred species-gene(s) associations have been repeatedly found (e.g. (Forsberg et al. 2014; Pradier and Bedhomme 2023). Studying the factors shaping and orienting horizontal transfers is key to understanding both bacteria genome evolution and the dynamics of horizontal dissemination of genes, in particular of antibiotic resistance genes. Indeed, resistance to antibiotics arises as an adaptive strategy, both in natural and clinical settings (Granato et al. 2019; Zhu et al. 2022) and can then spread, notably through the association with mobile genetic elements, to other bacterial populations and to new environmental niches. The strong selection for resistance evolution and dissemination incurred by the clinical use of antibiotics since the 1940s (Podolsky 2018) has led to an important burden imposed by antibiotic resistances and is increasingly referred to as a public health crisis (Murray et al. 2022).

Factors shaping and orienting horizontal transfers have been identified: a number of pre-HGT barriers or limitations explain why genetic material does not move freely (reviewed in (Popa and Dagan 2011; Acar Kirit, Lagator, and Bollback 2020; Burch et al. 2022). The most important of these factors are the requirement for donor and receiver to be in close physical proximity (usually through shared ecology), limited bacterial competency, mobile genetic element host range and resistance to bacteriophage (Thomas and Nielsen 2005). Once a gene has been transferred, its retention in the receiving genome and its vertical and horizontal propagation depends on the balance between the benefit it provides and the costs associated with its presence in the receiving genome, also called post-HGT barriers or limitations. The benefit is usually in terms of new function, for example the capacity to use new resources or to resist a drug or a heavy-metal (Chu et al. 2018; Woods et al. 2020) and often depends on the environment. As for the costs or limitations (reviewed in Baltrus 2013), they are first linked to the cost of carrying new genetic material, both in terms of replication of this material and in terms of perturbation of the transcriptome of the receiving cells, two phenomena which have been well documented for plasmid-mediated HGT (Harrison and Brockhurst 2012; San Millan et al. 2015). Second, some limitations come from the fact that the product of the introduced gene can interact negatively with other host proteins and perturb the host protein-protein interaction network (Burch et al. 2022). Finally, the introduced gene is beneficial only if it is efficiently transcribed and translated by the machinery of the receiving cell.

A key factor mediating the compatibility between a transferred gene and the host cell machinery is thought to be the differences in codon usage preferences (CUP) between the gene and the new host. Codon usage preferences refer to the preferential use of synonymous codons in coding DNA (Parvathy et al. 2022). CUP strongly differs between bacterial species and to a lesser extent between regions within a genome (Hershberg and Petrov 2008; Sharp and Li, 1987). Historically, CUP was thought to be the product of mutational bias and drift, with synonymous mutations seen as silent or neutral (Grantham et al. 1980). However, there are now numerous examples of non-neutral synonymous SNPs (e.g. (Lind et al. 2010; Bailey et al. 2014; Fragata et al. 2018; Lebeuf-Taylor et al. 2019; Bailey et al. 2021; Horton et al. 2021). Additionally, studies employing synonymous variants of a gene have highlighted how varying CUP results in different phenotypes and/or fitness (Kudla et al. 2009; Amorós-Moya et al. 2010; Cambray et al. 2018; Mittal et al. 2018; Walsh et al. 2020). The fitness effects of synonymous mutations and the role of selection in shaping CUP were initially interpreted as essentially being due to translational selection (Hershberg and Petrov 2008): codon usage frequencies are positively correlated with tRNA gene copy number, such that the codons which are preferentially used are codons for which the copy number of genes encoding the corresponding tRNA is the highest (Rocha 2004; Buchan et al. 2006). Horizontal transfer of a gene with CUP different from those of the receiving genome thus represents a break in the coevolution between CUP and the translation machinery, as the transferred gene is using many codons for which there are few tRNA to decode them in the receiving cells. These codons are rare codons in terms of CUP of the receiving genome. Use of rare codons is known to reduce translation efficiency and fidelity, leading to the production of a low quantity of functional protein and the accumulation of truncated and erroneous proteins, potentially toxic for the cell (Stoletzki and Eyre-Walker 2007; Drummond and Wilke 2010; Liu et al. 2021). More recently, the fitness effects of synonymous variation have also been linked to non-translation effects, either through its impact on mRNA secondary structure and transcript decay rate or through the presence of nucleotide motifs having a function besides that of encoding an amino acid sequence such as promoters, ribosomal binding sites and restriction sites (reviewed in Callens et al. 2021).

A horizontally transferred gene is likely to have CUP that differ from the receiving genome and in this context, the fitness effects of synonymous variation could be an important factor determining post-HGT success. Results from comparative genomic approaches support this as they show that HGT occurs more frequently amongst bacteria species with similar tRNA pools (Bahiri-Elitzur and Tuller 2021) and that accessory genes with CUP close to the core-genome tend to be retained more often in genomes (Callens et al. 2021). On the experimental side, most of the approaches investigating the phenotypic effects of synonymous variants (Kudla et al. 2009; Agashe et al. 2016; Mittal et al. 2018; Shaferman et al. 2023) find phenotypic differences between synonymous variants but they only use one model bacterial species, often being *Escherichia coli,* and do not include the across-species dimension of HGT. Additionally, in some of these studies (Kudla et al. 2009; Yan et al. 2016; Tian et al. 2017; Cambray et al. 2018; Mittal et al. 2018; Schmitz and Zhang 2021), the gene for which synonymous variants are generated is a fluorescent protein gene. Fluorescent protein genes are not expected to affect fitness and are not commonly transferred horizontally. Although they represent a powerful experimental model to quantify the amount of protein produced, they are not an ideal model to study the role of CUP in shaping and orienting HGT.

We set out to experimentally test the hypothesis that a mismatch between CUP of a transferred gene and that of the new host would determine the success of a HGT event and orient the transfers between species. To do so, the focal gene, for which synonymous variants were designed, was *aacC1*, a gentamicin resistance gene (Santucci and Krieger 2000). Besides public health relevance, antibiotic resistance genes represent a convenient model system as the phenotype they confer can be easily determined through classic resistance measures such as minimum inhibitory concentration (MIC) (Kowalska-Krochmal and Dudek-Wicher 2021) and selection pressure is easy to manipulate by varying antibiotic concentrations in the environment. Our biological system was composed of three bacterial species *Acinetobacter baylyi*, *E. coli* and *Pseudomonas aeruginosa,* with very contrasted CUP (Table S1). The strains we used are each closely related to pathogenic bacteria species or strains (Huai et al. 2019) which together make up a large proportion of deaths attributed to or associated with antibiotic resistance (Murray et al. 2022).

We hypothesized that variants which better matched the host’s CUP would provide higher resistance. By introducing our *aacC1* synonymous variant collection into the three chosen bacteria species, we established that they indeed conferred very different levels of resistance to gentamicin within each species. Additionally, the overall mean resistance differed from one species to another and variant’s resistances did not rank in identical order across species, revealing an interaction between gene variant and host genome in determining the phenotype. However, in contradiction to our original hypothesis, the average similarity of codon usage between the transferred gene and the host genome was not the main determinant of resistance level for each species-variant combination. The role of local codon usage and internal motifs were then further investigated. Species-specific effects were revealed which highlight both the benefit of our multi species approach and the difficulty of predicting resistance levels and HGT success, from nucleotide sequences.

### Results Section

The basic premise of our experimental approach was to determine the impact of codon usage preferences on the functional success of horizontal transfer by mimicking the transfer of synonymous versions of an antibiotic resistance gene between three bacteria species. The experimental system was composed of the *aacC1* gene, as the resistance gene, and *E. coli*, *P. aeruginosa* and *A. baylyi,* as the bacteria species in which the synonymous versions were transferred. *aacC1* produces an aminoglycoside-acetyltransferase which provides resistance to gentamicin by transferring an acetyl group to the antibiotic thus impeding drug fixation on its target, the 30S ribosomal subunit. *P. aeruginosa* favors GC rich synonymous codons while *A. baylyi* favors GC poor codons. *E. coli* does not favor either, in line with its overall genome GC content of approximately 50%.

### Design of synonymous gene variants to match and mismatch the codon usage preferences of three bacterial host species

We designed synonymous variants of the *aacC1* gene by varying codon positions 20 to 183 (Figure 1A, 31-549 bp). The amino acid sequence of each variant remained identical to the wild-type *aacC1* sequence (GenBank: AAB20441.1). We specially designed this variable region to match the codon usage of each of the bacterial species (Figure 1C, S1). To design the variable region, we used the COUSIN analysis tool (Bourret et al. 2019): for each variant and each amino acid position independently, one of the synonymous codons was assigned in a semi-random manner, based on the frequencies of use in the codon usage table of one of the host species. Out of the 90 variants designed, 31 were retained (see methods) including a variant in which the most frequently used synonymous codon was attributed for each amino acid for each species.

**Figure 1.**
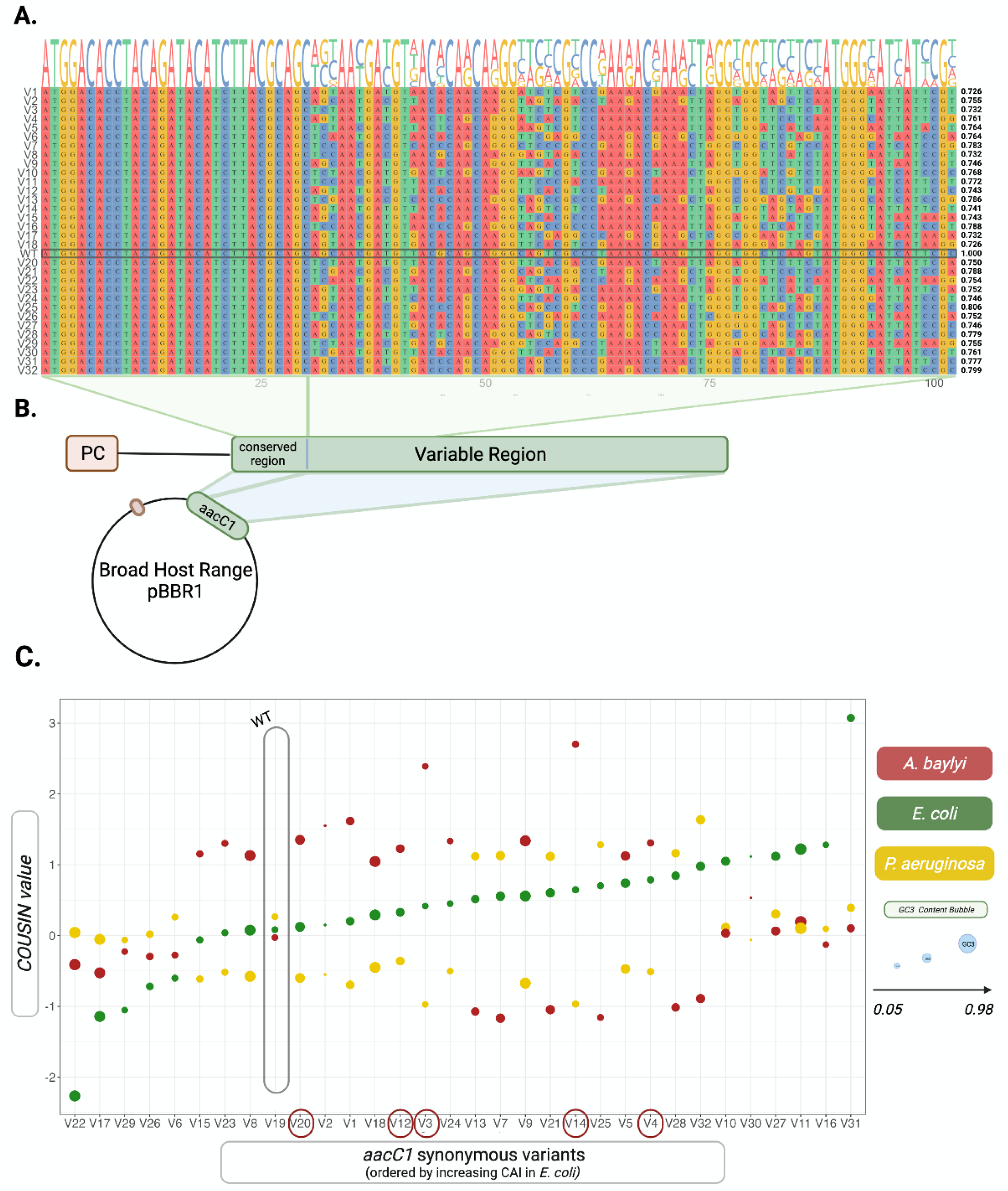
*aacC1* synonymous variant design and constructs. (A) First 34 codons of the Synonymous variant sequences. The first 10 codons (30bps) were conserved across variants. The remaining codons were assigned different synonymous codons in semi-random manner, guided by the codon usage of the bacteria species used. Pairwise identity with wt *aacC1* is shown to the right of each sequence. (B) Variant constructs were composed of an integron derived promoter and 5’ UTR, followed by the *aacC1* gene and cloned into a pBBR1 broad host range plasmid. Variant plasmid sequences differed only by the variable region within *aacC1*. (C) Codon usage similarity (COUSIN) for the 32 synonymous *aacC1* variants in the three host-species used. Variants are ordered by increasing COUSIN values in *E. coli*. GC content at the third base of each codon (GC3) is represented by bubble size. The five variants that could not be transformed into *A. baylyi* are circled in red on the x-axis.

In order to minimize the effects of differences in translation initiation around the translation start site due to mRNA folding, we conserved the first 30 bps of the *aacC1* sequence. Recent studies (Liu 2020; Cambray et al. 2018; Liu et al. 2021) suggest that synonymous variation at the 5’ end of the coding sequence is responsible for the lion’s share of phenotypic differences between synonymous variants, through the effect of the nucleotide sequence on mRNA folding energy and consequently on ribosome accessibility to the Ribosomal Binding Site. Thus, we choose to conserve this region in order to untangle the effects of codon usage preferences on translation initiation and on translation efficiency and accuracy. The stop codon was also identical for all variants.

The 31 synonymous variants (and the wild-type sequence) of the *accC1* gene covered a wide range of codon usage match within each species and presented important shifts in codon usage match from one species to another (figure 1C, S1). The level of codon usage match was quantified with COUSIN (COdon Usage Similarity INdex), an index which measures the similarity of codon usage of a focal sequence to a set of reference genes normalized to a Null Hypothesis of equal usage of synonymous codons. COUSIN and the classical Codon Adaptation Index (CAI, Sharp and Li 1987) are positively correlated (Figure S2) but by normalizing to equal usage of synonymous codons, COUSIN allows for direct comparison between genomes (Bourret et al. 2019). The 32 variants also differed in GC content (figure 1C), and had pairwise identities to the wt ranging from 0.726 to 0.806 (median = 0.755) (Figure 1A). The pairwise identity between variants ranged from 0.621 to 0.940 (median = 0.759) (Figure S3). The third codon GC content (GC3) of the variants ranged from 5.98% to 98.37% (median = 44.57%, wt = 59.78%).

The 32 variants were synthesized and cloned into a natural broad host range plasmid downstream of a natural integron-derived promoter and 5’UTR region (Figure 1B). The 32 plasmid versions, each carrying one synonymous variant, were successfully introduced in *E. coli* and *P. aeruginosa* by electroporation. 27 of the variants were introduced into *A. baylyi* by natural transformation. The remaining five variants could not be transformed (Figure 1C). These non-transformed variants show high COUSIN values in *A. baylyi* compared to the 27 transformed variants (mean COUSIN value of the five variants = 1.80, mean COUSIN of the transformed 27 variants = 0.24). The 5 variants also have much lower GC3 content (mean GC3 of the five variants = 19.13%, mean GC3 of the transformed 27 variants = 50.58%).

### Synonymous variation modulates resistance levels in three different bacterial hosts

Resistance levels were determined by growing all *aacC1* synonymous variant * species combinations across a range of gentamicin concentrations. We chose to measure resistance phenotype, as opposed to transcript or protein levels, in order to measure the overall impact of our synonymous variants on the host cell. We used two variables to represent resistance to gentamicin in our variants. The first, minimum inhibitory concentration (MIC) is defined here as the lowest concentration at which the variant does not reach an OD 600 value of 0.1 (see methods). The second measure, half maximal inhibitory concentration (IC50), is the concentration of antibiotic at which the growth in the absence of antibiotic is reduced by 50%. Here, IC50 is calculated using the Area Under the Curve (AUC) for 24 hours (or 48 hrs for *A. baylyi*) calculated from growth curve measures at serially diluted concentrations of gentamicin. The AUC growth metric was chosen as it integrates information about lag time, growth rate and carrying capacity (Ram et al. 2019). AUC, lag time, and OD max (carrying capacity) measures for each variant in each gentamicin concentration are available in the supplementary materials (Figures S4-S6).

We found a strong statistically-significant interaction between species and variants for IC50 AUC (two-way ANOVA, F = 4.22, *p* = 5.84e^-14^). Species and variants had significant effects (species, F = 91.853, *p* = <2e-16; variant F = 2.575, *p* = 3.15e-05). Mean IC50 ranged from 20µg/mL gentamicin in *A. baylyi to* 1024 µg/mL in *P. aeruginosa* (Figure 2A). The variant effect on IC50 AUC was also significant within each species (Table S2): the modulatory capacity of synonymous variation on resistance levels held across all three bacterial species, as illustrated in Figure 2A, with IC50 differing by a 487.26, 7,83 and 6,66 fold in *A. baylyi*, *E. coli* and *P. aeruginosa*, respectively. MIC values ranged in *P. aeruginosa* from 800µg/mL to >25600 µg/mL, whilst they ranged from 25µg/mL to >25600µg/mL and 6.25µg/mL to >400µg/mL in *E. coli* and *A. baylyi*, respectively (Figure S7). “Variant” had a statistically significant effect on MIC within each of the species (Table S2).

**Figure 2.**
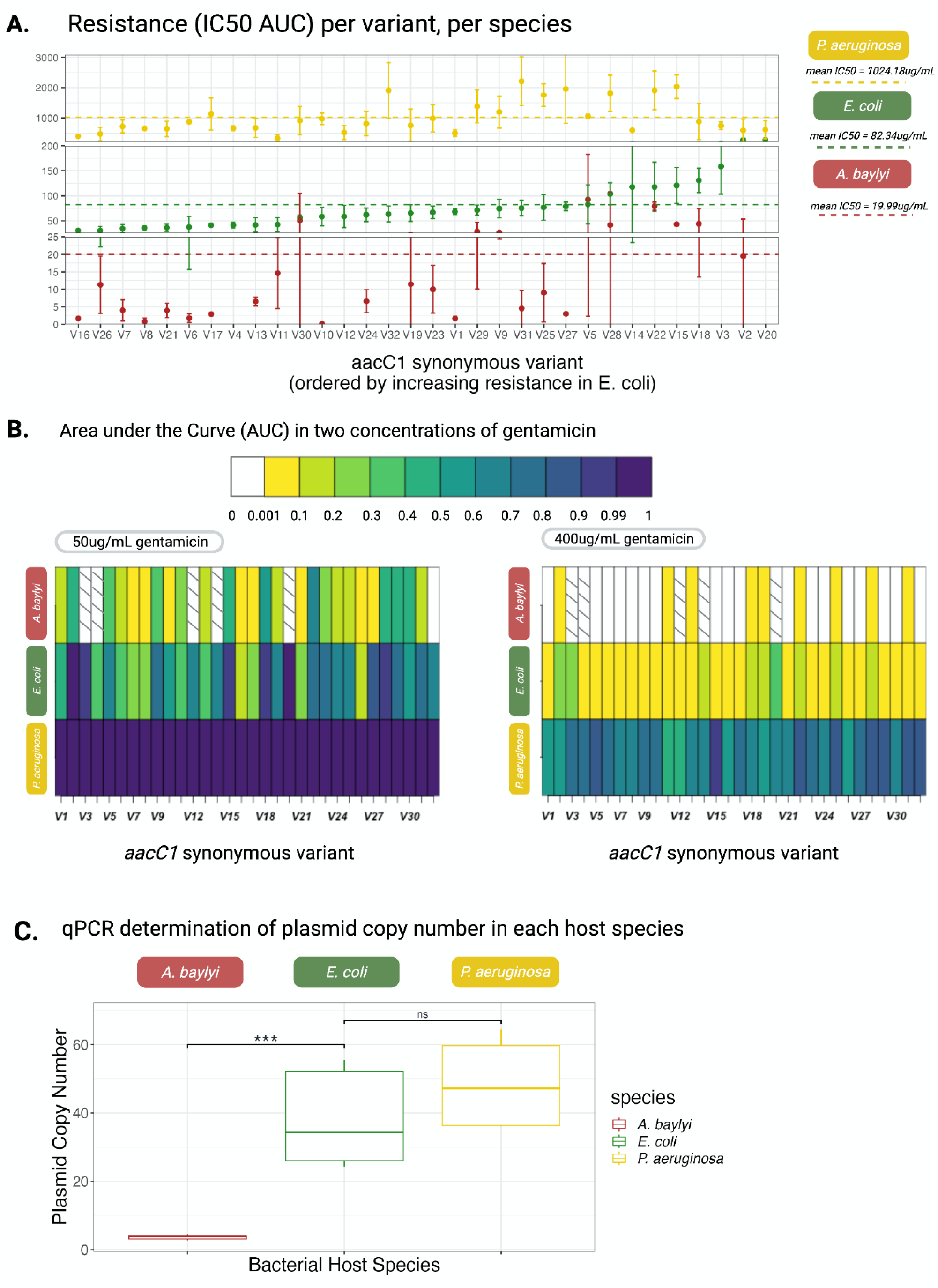
Synonymous variants confer different resistance levels. (A) Resistance levels are represented by IC50 AUC *i.e.* the concentration of gentamicin required to reduce the bacterial growth by 50% compared to growth in the absence of gentamicin. Synonymous variants are ordered on the x-axis by increasing IC50 AUC in *E. coli*. Mean IC50 for each species is represented by a coloured dashed line. (B) AUC of the three bacteria species carrying the *aacC1* variants in 50µg/mL and 400µg/mL of gentamicin. AUC is represented as relative to AUC in 0µg/mL gentamicin. A white box indicates that the variant-species combination was not tested in this concentration but this concentration is above the MIC for this variant and it can thus be confidently inferred that the AUC is null. Dashed boxes indicate the five variants that could not be transformed in *A. baylyi*. Values represented by the darkest purple colour were set to 1. These variant-species combinations were not tested in this concentration but had a relative growth of >0.9 in a lower concentration of gentamicin. (C) Plasmid copy number, quantified by qPCR, in each host variant. Significant differences in plasmid copy number are indicated by ***, non significant are indicated by ns.

Given the strong species effect (Figure 2A), we sought to determine if differences in resistance levels were due, in part, to differences in plasmid copy number (PCN). We quantified PCN, the number of plasmid copies in each bacterial species, by qPCR and observed significant differences (One way ANOVA, F = 51.8, *p* = 1.96e-09, Table S3) in copy number between species (PCN: *A. baylyi* 3.67+/-0.65, *E. coli* 38.00+/-12.60, *P. aeruginosa* 48.72+/-11.37). Notably, PCN ranked in the same order as the average resistance level in each species. The lower copy numbers observed in *A. baylyi* (compared to *P. aeruginosa* and *E. coli*) could explain the overall lower average resistance levels in this species (Figure 2A, 2C).

The combination of the species and synonymous variant effects in determining the resistance level is very likely to affect the immediate success of HGT, as illustrated in Figure 2B: the AUC at a given gentamicin concentration differs strongly depending on the receiving species and the received synonymous variant. In the 50µg/mL environment, all of the *P. aeruginosa* grow as well as in the absence of antibiotics but when the variants are transferred into *E. coli*, there is a large range of AUC levels (from 0.1 to 1), showing that some variants permit growth and others have a clear disadvantage post-HGT. This effect is even more pronounced in *A. baylyi*, suggesting that many of the variants, if transferred into a 50µg/mL environment would not succeed post-HGT and be selected against. At 400µg/mL, *P. aeruginosa* variants grow at different rates (AUC range from 10.48 to 23.06), influencing their post-HGT success in the environment. Inversely, almost all of the variants, when transferred into *E. coli* and *A. baylyi,* show very little growth, preventing post-HGT success in 400µg/mL gentamicin. The significant “variant” effect actually holds in most individual gentamicin concentrations for AUC, lag and OD max (One-way ANOVA, Table S4).

### Similarity with host codon usage preferences is not a strong determinant of variant resistance phenotype

The Codon Adaptation Index (CAI) was first described by (Sharp and Li, 1987) and linked the biased use of synonymous codons in highly expressed genes to their gene expression, putting forward the hypothesis that codon usage bias (CUB) is correlated with expression and thus potentially under selection. Since then, a multitude of codon usage indexes have been developed to piece apart the relationship between CUB and phenotype, through their impact on transcription, translation, mRNA stability and co-translational folding (reviewed in Bahiri-Elitzur and Tuller 2021).

We sought to determine if the observed differences in resistance phenotype could be explained by the effect of CUP on translation efficiency and fidelity. To do so, we used different codon usage indexes: CAI (Sharp and Li, 1987), COUSIN (Bourret et al 2019) and also tAI which measures the match between the CUP of a sequence and the tRNA gene pool of the host genome (dos Reis et al. 2003). The last index used was |Δ(GC3)|, measuring the absolute value of the difference in GC3 between a sequence and the host genome.

A significant positive correlation was found between COUSIN and resistance levels in *P. aeruginosa* (r=0.35, p=0.049) but not in the other two species (figure 3B). No significant correlation existed between CAI or tAI and resistance level in either of the species (Figure 3). We also find a significant positive relationship between |Δ(GC3)| and resistance in *E. coli* (r=0.38, p=0.032). We further explored correlations between codon usage indices and AUC within individual gentamicin concentrations (Table S5). While IC50 measures allow us to determine resistance levels for each variant, individual concentrations can be regarded as unique environments in which a variant may be horizontally transferred. In agreement with the inter-concentration effect, we observed significant correlations between AUC and COUSIN in *P. aeruginosa* at high gentamicin concentrations and between AUC and |Δ(GC3)| in *E. coli* at intermediate concentrations. Additionally, we observed a number of significant correlations in individual gentamicin concentrations that were masked in the IC50 results, such as positive correlations between AUC and CAI in *P. aeruginosa* and between AUC and |Δ(GC3)| in *A. baylyi* and a negative correlation between AUC and |Δ(GC3)| in *P. aeruginosa* for high gentamicin concentrations.

**Figure 3.**
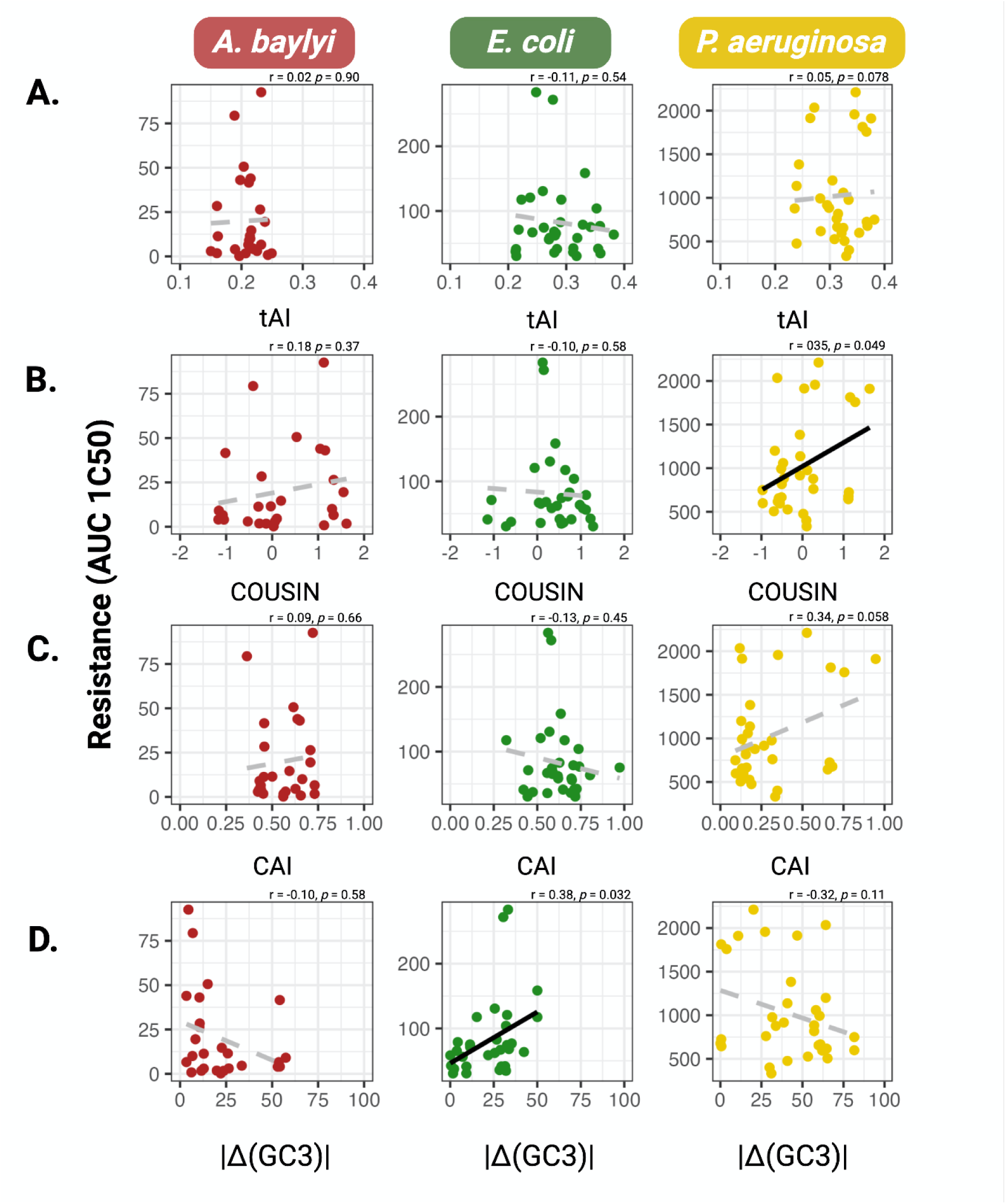
Codon Usage Preferences and Resistance. (A) Pearson correlation between resistance (IC50 AUC) and tAI. Significant correlations (p < 0.05) are represented by a black regression line. Non-significant Pearson correlations are represented by a grey dotted regression line. (B) Pearson correlation between resistance (IC50 AUC) and COUSIN. (C) Pearson correlation between resistance (IC50 AUC) and CAI. (D) Pearson correlation between resistance (IC50 AUC) and |Δ(GC3)|.

The results presented here provide some insight into the mechanisms determining the modulatory effect of synonymous variation on resistance levels and bacterial growth and fitness in the presence of antibiotics. However, codon usage similarity averaged over the gene only partially explains the phenotypic differences between synonymous variants in two out of the three species. This prompted us to explore the relationship between resistance and local codon usage.

### Localized sequences within the *aacC1* gene partly modulate resistance phenotype

Overall similarity in CUP to the host genome could not fully explain the large variation of resistance levels amongst the *aacC1* variants. But it is possible that the effects are due to the codon usage of key positions or segments of the variants and that these effects are not captured when using CUP indices calculated over the whole variant sequence. To try and reveal these potential local codon usage effects, we calculated CAI, COUSIN, tAI and |Δ(GC3)| values for 30bp sliding windows across the *aacC1* gene, with 15bp steps. This window size was chosen as it represents the ribosome footprint (Martens et al. 2015). The ribosome footprint, or rather the 10 codons (30 bp sequence) covered by an individual ribosome is predicted to influence translation elongation rates and translational pauses. Translational elongation rates and pausing in turn influence protein folding and possibly translation abortion rates (Verma et al. 2019), modulating protein levels and protein quality and thus potentially protein function, phenotype and fitness.

We then looked for correlations between variant resistance levels (IC50 AUC) and the codon index values of individual 30bp windows. Unsurprisingly, we captured significant correlations in sliding windows both for *P. aeruginosa* between COUSIN and resistance (15/25 sliding windows, Figure 4D) and in *E. coli* between |Δ(GC3)| and resistance (16/35 sliding windows, Figure 4C), consistent with the results obtained with codon usage index averaged across the whole gene. In *P. aeruginosa,* the significant positive correlations with COUSIN values were localised in the regions 15-90 bp and 450-525 bp (Figure 4D). In *E. coli*, the significant negative correlation with |Δ(GC3)| was distributed across the sequence (Figure 4C).

**Figure 4.**
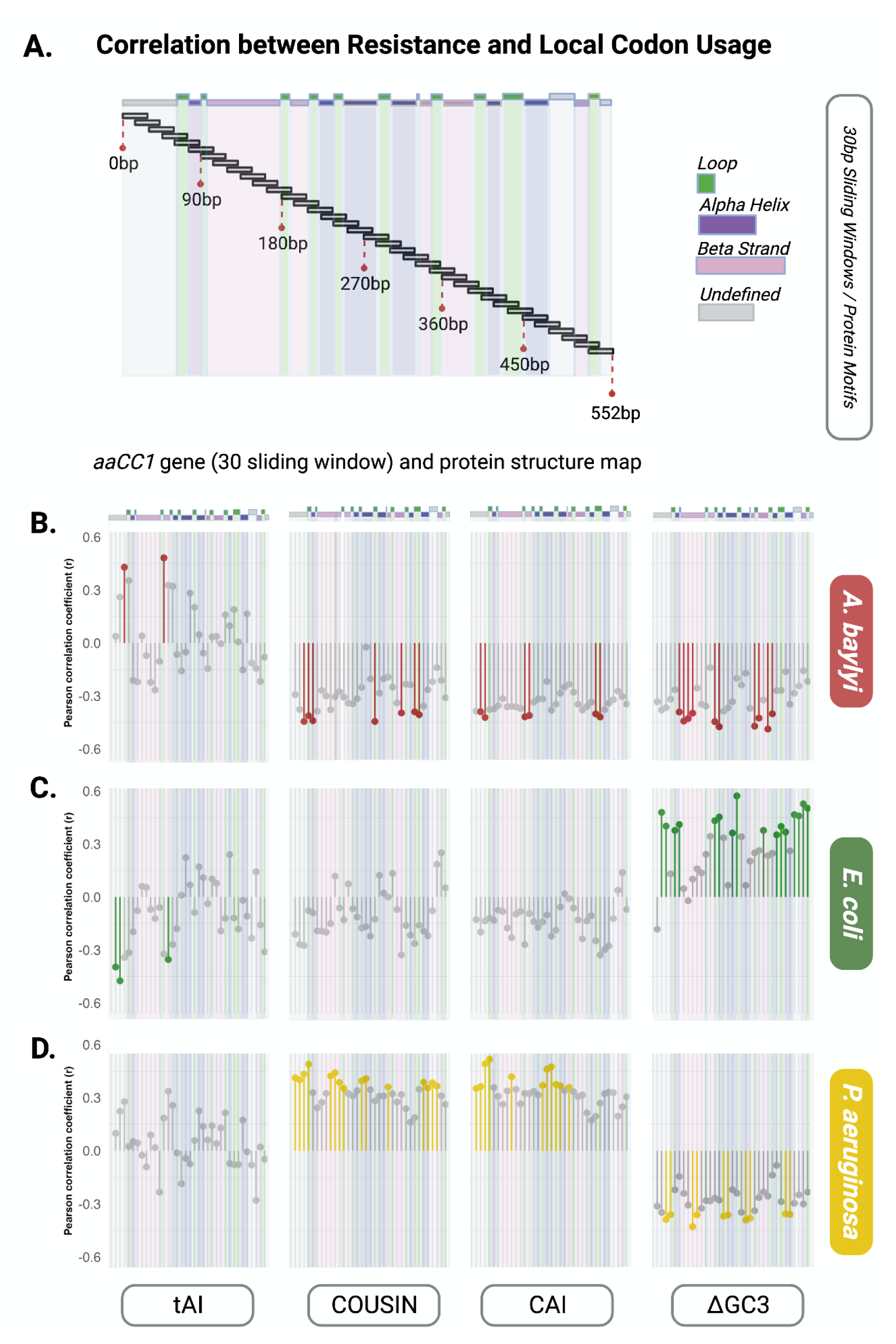
Local codon usage and resistance. (A) Visual representation of the 36 30 bp-sliding windows of the *aacC1* gene. Protein structural elements, *i.e.* alpha helices, beta sheets and loops are represented in purple, pink and green, respectively. Undefined elements are represented in grey. (B) Pearson correlation coefficients between resistance (IC50 AUC) and tAI, COUSIN, CAI or |Δ(GC3)| in *A. baylyi*. Lollipop graph icons represent the 35 r value for each sliding window. Note that window 0-30bp is common to all *aacC1* variants and is thus excluded from correlation tests. Non-significant correlations (p-value >0.05) are represented in grey. (C) Pearson correlation coefficients between resistance (IC50 AUC) and tAI, COUSIN, CAI or |Δ(GC3)| in *E. coli*. (D) Pearson correlation coefficients between resistance (IC50 AUC) and tAI, COUSIN, CAI or |Δ(GC3)| in *P. aeruginosa*. All correlation test values, including the Benjamini-Hochberg (BH) corrections not included here, are available in the Tables S6 and S7. Supplementary Figure S4 mirrors Figure 4 but includes BH corrections.

COUSIN shows significant negative correlation with resistance in *A. baylyi* (7/35 sliding windows, Figure 4B). No such significant correlation was observed in *E. coli* (Figure 4C). In *A. baylyi,* this significant negative correlation was localised at the beginning of the sequence 45-105 bp and at 420-465 bp (Figure 4B), similar to the location in *P. aeruginosa*.

For |Δ(GC3)| content, we observed significant negative correlations with resistance in *A. baylyi* (10/35 sliding windows, Figure 4B). In *P. aeruginosa*, we observed significant positive correlations (10/35 sliding windows, Figure 4D). In *A. baylyi* this resistance - |Δ(GC3)| negative correlations are localised from 30-120 bp, from 270-315 bp and from 420-480bp (Figure 4B). In *P. aeruginosa*, the positive correlations are from 135-195bp, from 255-375 bp and from 450-540 bp (Figure 4D).

For CAI, significant negative correlations with resistance were observed in *A. baylyi* (6/35 sliding windows) and significant positive correlations were observed in *P. aeruginosa (*11/35 sliding windows), mirroring the results for COUSIN (Figure 4C). No significant correlation was observed in *E. coli.* In *A. baylyi*, this resistance vs CAI negative correlations are localised from 15-105 bp, 180-225 bp and from 420-465 bp. In *P. aeruginosa*, the positive correlation was from 15-105bp and from 240-360 bp (Figure 4C).

For tAI, there was no significant correlation in *P. aeruginosa* (Figure 4D). In *A.baylyi,* two sliding windows, 45-75 bp and 180-210 bp, correlated positively with resistance, (Figure 4B). In *E. coli*, resistance correlated negatively with tAI in three windows (15-60 bp and 195-225 bp) (Figure 4C).

The significant correlations presented in Figure 4 are based on uncorrected *p*-values. Corrected *p-*values, using the Benjamini-Hochberg (BH) false discovery rate correction method are available in Figure S8 and Table S6. The false discovery rate (FDR) chosen for this experiment was 0.25, with the acceptance that 1 in 4 significant correlations could be false positives. Thus, 1 in 4 significant correlations in Figure S8 could be false positives, with the BH correction increasing the number of significant tests compared to Figure 4. Note that the 15bp overlap between sliding windows means that our individual tests are not 100% independent. Additionally, given that the majority of tests had a *p-*value <0.25 (Table S7) and the co-locations of significant results along the *aacC1* sequence, we are confident in the general trends observed in our results (see Mcdonald J.H. 2014).

The local effects detected are thus not located on the same segment of the gene in the three species nor across the different indices and they do not seem to correspond to secondary structure of the protein (figure 4E). This was further confirmed by the absence of correlation between the codon usage indices and the resistance level for the concatenation of the segments belonging to each of the four secondary structure classes (see table S8).

### Translational bottlenecks do not explain differences in resistance levels between synonymous variants

Another way through which local codon usage could affect the level of resistance is the presence, length and position of translational bottlenecks. To test this hypothesis, we have used two ways of defining and quantifying translational bottlenecks. First, following Navon and Pilpel (2011), we identified translational bottlenecks as the 30bp window with the lowest tAI. Navon and Pilpel (2011) have shown, in *E. coli,* that high bottleneck strength and bottleneck proximity to the start codon are positively correlated to protein production. However, we found no significant correlation between bottleneck position or strength and resistance levels in the three host species (Figure 5B, 5C).

**Figure 5.**
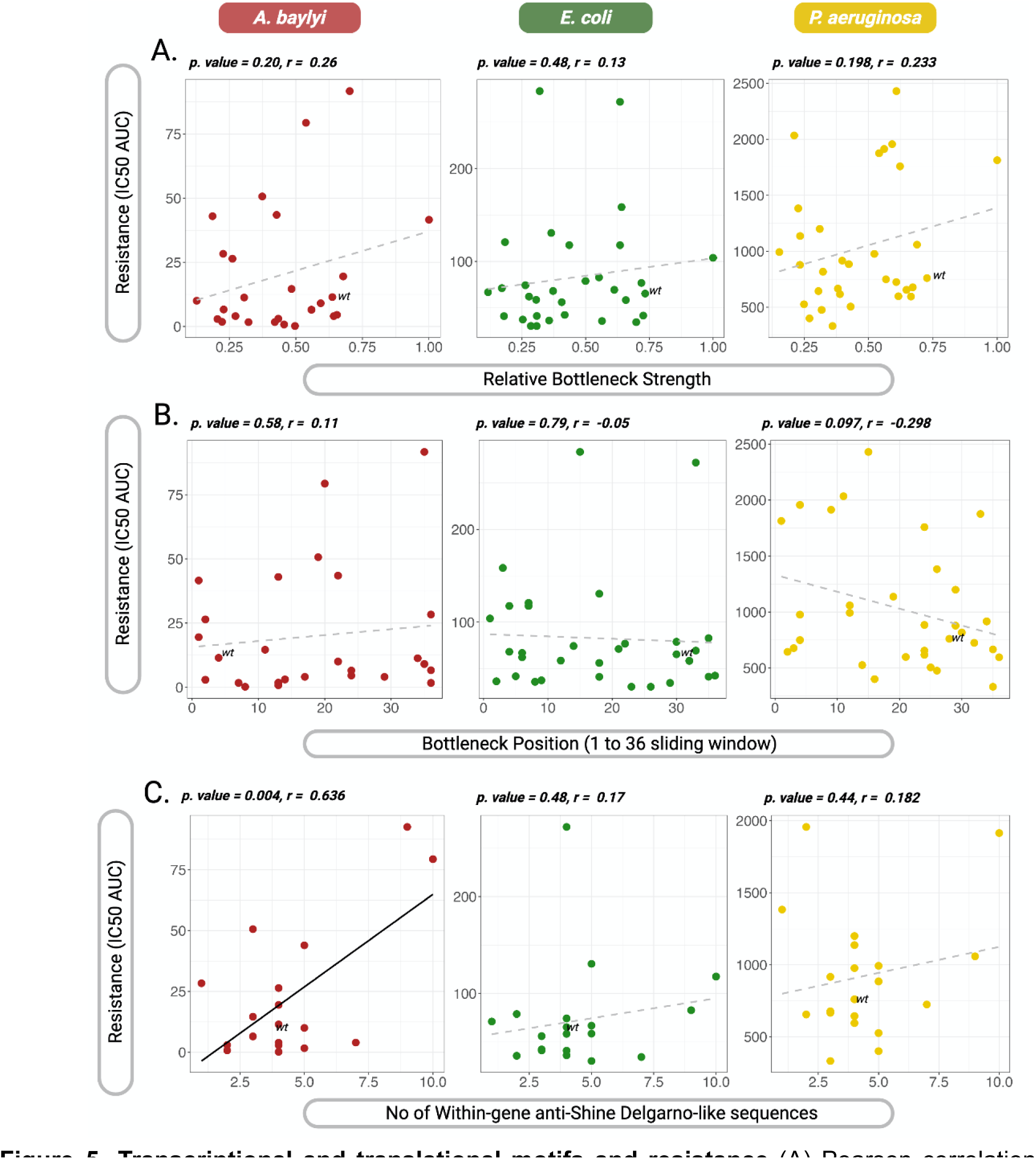
Transcriptional and translational motifs and resistance. (A) Pearson correlation between resistance (IC50 AUC) and relative bottleneck strength. Relative bottleneck strength is the minimum tAI value divided by the mean tAI value of the 36 sliding windows (B) Pearson correlation between resistance (IC50 AUC) and bottleneck position. (C) Pearson correlation between resistance (IC50 AUC) and the number of hexamers within the *aacC1* variant with < -7 kcal/mol binding free energy (strong anti-SD-like sequence).Significant correlations (p < 0.05) are represented by a black regression line. Non significant Pearson correlations are represented by a grey dotted regression line.

Second, we tested for a correlation between resistance and the number of rare codon chains. We define rare codons for each species as codons that are underrepresented in the host CUT. More specifically, we defined rare codons as those with less than half their expected frequency under the null hypothesis of equal frequency within a synonymous codon family (e.g. for a codon within a four synonymous codon family, a rare codon is a codon with a frequency <0.125). We hypothesized that a high number of rare codons and high frequencies of rare codon chains would produce translation bottlenecks slowing down translation, increasing translation errors and potentially leading to ribosome abortions which can occur during prolonged translational pausing. We tested for correlations between the number of rare codons and resistance levels and also between the number of chains of two, three, four and five rare codons and resistance levels. We did not find any significant relationship in any of the species (Table S9).

### Internal Shine-Dalgarno-like motifs and secondary mRNA structure do not explain differences in resistance levels between synonymous variants

The Shine-Dalgarno (SD) sequence, or bacterial ribosome binding site (RBS), is complementary to the 3’ end of the 16S rRNA of the ribosome and controls translation initiation. Within gene SD or SD-like sequences are expected to have deleterious effects because they can cause within-gene translation initiation and generate ribosome conflicts. Their presence has been shown to impose selective constraints on the surrounding sequences (Hockenberry et al. 2018). It is possible that our semi-random design of synonymous variants introduced within-gene SD-like sequences and we hypothesized that there would be a negative relationship between resistance and a high frequency of within-gene SD-like sequences. To test this, we looked for a potential relationship between the number of hexamers with a strong affinity to the anti-SD sequence and the resistance level. We found no significant relationship in either *E. coli* or *P. aeruginosa* and a significant positive correlation between the number anti-SD-like hexamers (< -4 kcal/mol free binding energy) (Hockenberry et al. 2018) and resistance in *A. baylyi* (Figure 5A, Table S10). This significant positive relationship remained when tested in strong anti-SD-like sequences (< -7 kcal/mol binding free energy, Table S10). This result being contrary to our prediction, we looked for a potential explanation: the *aacC1* wild-type sequence used, initially found in *Serratia marcescens,* shares almost 100% sequence alignment with a shorter, *P. aeruginosa* derived *aacC1* variant, bar the first 69 bps. Thus, the first 87 bp (includes the 18bp AU1 epitope following ATG of our variants) are not required for gentamicin acetyltransferase activity and the shorter protein variant could potentially confer resistance more efficiently than our *aacC1* variant. We thus split the *accC1* gene into two segments and correlated resistance and anti-SD-like sequence frequency separately for the beginning 87 bp segment and the end 465 bp segment (required for enzymatic activity). We hypothesized that the positive correlation between resistance and anti-SD-like frequency was mainly due to the 87 bp fragment. Surprisingly, the significant positive correlation we observed in *A. baylyi* in the full sequence was lost in the 87 bp segment but maintained in the remaining 465 bp segment (Table S11).

Finally, the *aacC1* variants were identical for the first 30 bps to limit differences between variants in the 5’ end secondary structure of the mRNA and the associated effects on translation initiation. However, previous studies (e.g. Cambray *et al*. 2018) identified an effect of mRNA structure of a longer mRNA stretch. We tested whether this was also the case in our synonymous variants by looking at the correlation between mRNA folding energy in the 90 bps around the start codon (−30:60 bp) and resistance. We found no significant correlation in any of the three species (Table S11).

## Discussion

We have shown that synonymous variants of a gentamicin resistance gene confer strongly different resistance levels in three distant bacteria species *A. baylyi*, *E. coli* and *P. aeruginosa*. Our resistance measures show a strong species effect, explained in part by differences in plasmid copy number. Additionally, there is an important variant * species interaction, with the variants conferring the highest resistance being different from one species to another. This species-specific effect of synonymous variation points to codon composition as a factor influencing immediate post-HGT success and orienting HGT. However, the average similarity in codon usage between a synonymous variant and the genome of the recipient species only explains a limited part of the variation in resistance levels in one of the species, *P. aeruginosa*.

The strongest determinant of the resistance conferred by the synonymous variants of the *aacC1* gene was the species in which it was introduced. The copy number of the broad-host range plasmid pBBR1, used as a vector for the synonymous variants, strongly differs between the three host species used. The three species rank in the same order for PCN and average gentamicin resistance level conferred, indicating that PCN might explain the differences in average resistance level between species through a resistance gene dosage effect. Different mechanisms of PCN regulation have been described (Del Solar and Espinosa 2000; Nordström 2006), but they all involve only plasmid sequences coding for negative regulation mechanisms, such as antisense-RNA. Previous studies have linked chromosomal point mutations and PCN in E. *coli* (Lopilato et al. 1986; Chiang et al. 1991; Xu et al. 1993; Yang and Polisky 1999; Ederth, et al. 2002) but the interspecific variation in plasmid copy number is not broadly documented and the mechanism behind it are not known. Interestingly though, (Pena-Gonzalez et al. 2018) report copy number variation for two plasmids across *B. cereus* and *B. anthracis* strains with no phylogenetic signal in the PCN variation but a positive correlation between the copy number of the two plasmids, suggesting a host-driven copy number control common for the two plasmids.

Very recently, (Alonso-del Valle et al. 2023) documented copy number differences for the same antibiotic resistance plasmid between species (*E. coli* and three Klebsiella species) and between clinically relevant strains within species. Additionally, they report a positive correlation between PCN and the level of resistance conferred. All together, these results strongly suggest that PCN variation across strains and species has implications for immediate HGT success through gene dosage, as genes transferred horizontally on plasmids will produce different phenotypes depending on the PCN in the new host. Additionally, it is possible that the transcription level from a same promoter differ from one species to another: by transferring a library of regulatory elements (RE) into three bacteria species (*Bacillus subtilis*, *E. coli* and *P. aeruginosa*), Gomes et al. (2020) show a generally lower promoter activity for a same RE in GC-poor species. Interestingly, out of the three species they use, two are in common with our experimental system and the third (*Bacillus subtilis*) has a GC content similar to the one of our third species, *A. baylyi*. Their results suggest that the expression of *aacC1* would be higher in *P. aeruginosa* than in *E. coli* and higher in *E. coli* than in *A. baylyi*, an effect going exactly in the same direction as the PCN effect. The modeling approach in Alonso-del Valle et al. (2023) established that in their system, the level of resistance conferred to a strain was a strong determinant of the resistance gene-strain association and of the community composition. In the longer term, PCN is also known to influence the probability of segregational loss, and so plasmid stability, as well as the evolutionary dynamics of the genes carried (Rodriguez-Beltran et al. 2018). Strain- and species-specific PCN and expression level is thus very likely to be an important factor in determining the post-HGT success and evolution and in orienting HGT over the long term. Understanding the host PCN regulation mechanisms as well as host-specific expression levels is thus important for a better understanding and prediction of antibiotic resistance propagation.

The differences in resistance conferred by our synonymous variants within each of the three species used illustrate the potential of synonymous variation to modulate the phenotype likely through the expression level of genes and the functionality of the proteins they encode. Our results are in line with previous studies, which have shown synonymous variation impact mRNA and protein quantities (Kudla et al. 2009; Agashe et al. 2013), mRNA toxicity levels (Mittal et al. 2018), protein folding (Walsh et al. 2020), enzymatic activity levels or protein functionality (Amorós-Moya et al. 2010; Agashe et al. 2013; Yannai et al. 2018) and lead to differences in growth rates or fitness (Agashe et al. 2013; Mittal et al. 2018; Yannai, Katz, and Hershberg 2018; Shaferman et al. 2023). Taken together, they show that synonymous variation affects cell function on multiple levels from mRNA levels, to protein levels and protein folding, producing measurable fitness effects. However, all previous studies have been conducted within one species, *E. coli* for a majority of them, and do not provide information of the species-specificities of these effects and do not test directly the implications for HGT. Our approach, by introducing the same collection of synonymous variants in three distant species with contrasted codon usage allowed both to tease apart species-specific effects of synonymous variation and to mimic the horizontal transfer of a same variant across species.

Phenotypic characterisation of the *aacC1* variants actually showed that the relative levels of resistance conferred by each synonymous variant were not conserved across species. These species-specific phenotypic effects indicate differing compatibility between the transferred variant genes and the receiver bacterial genomes, with individual variants being more compatible in one genome and less compatible in another. This compatibility has clear implications for horizontal gene transfer outcomes. For example, a variant which confers a certain level of resistance in one species and provides a fitness advantage in the presence of antibiotics, may potentially be transferred into other species in the bacterial community. However, if the variant is incompatible or less compatible with the new host species, the resistance phenotype will not manifest and no fitness advantage will be conferred, preventing the retention of the transferred gene and its further propagation.

A potential mechanism determining the compatibility of a synonymous variant to a new host species is the similarity in codon usage between the variant and the host genome. Hence, we originally hypothesized that variants would differ in translation efficiency and accuracy, resulting in resistance levels that correlated with codon usage similarity indices. Interestingly, the resistance conferred by a variant in a species did not generally correlate with average codon usage similarity indices. We did, however, find a significant correlation between resistance and COUSIN in *P. aeruginosa* and between resistance and |Δ(GC3)| content in *E. coli* (Figure 3). We further reasoned that the synonymous variants differed by a large number of positions and that the global effect might not be due to the sum of small individual effects but due to the effects of a limited number of these positions. Otherwise said, codon usage might have localised effects that become diluted and thus masked when considering codon usage indices averaged over the whole gene sequence. To explore this possibility, we examined the correlation between resistance and localized codon usage within the *aacC1* gene, in different ways.

First, we correlated our resistance levels with codon usage indices calculated over 30bp sliding windows and identified significant positive correlations between resistance levels and COUSIN in *P. aeruginosa* and between resistance levels and |Δ(GC3)| in *A. baylyi* for some windows, mirroring the results at the whole gene scale. Additionally, some windows showed positive correlations between CAI and resistance level in *P. aeruginosa* and negative correlations between resistance and CAI and COUSIN in *A. baylyi* and between resistance and |Δ(GC3)| in *P. aeruginosa* (Figure 4). These results validate the idea that there are localised codon usage effects that could not be identified using indices averaged over the whole sequence. The localised effects identified are not in the same direction and for the same index across species and do not concern the same parts of the protein. This suggests a species specificity in terms of sensitivity to local codon usage similarity. Additionally, there is no clear correspondence between sections of the gene that show local codon usage sensitivity and the secondary structure of the protein, making it difficult to generate further hypotheses about the mechanistic link (translation speed, translation fidelity, protein folding) between local codon usage and resistance level. A second way through which local codon usage could have an effect on translation is by the presence of rare codon bottlenecks. Consecutive rare codons have indeed been shown to increase translation errors and ribosome stalling and drop-off (Aguirre et al. 2011; Mordret et al. 2019). However, rare codon chains of different length and translation bottlenecks did not explain the resistance conferred by the *aacC1* synonymous variants in any of the three species used.

Finally, we explored the potential role of the presence of sequence motifs within the synonymous variants coding for information beyond the amino acid sequence, whose function and activity could interfere with transcription or translation of the synonymous variant (reviewed in Callens et al. 2021). In particular, the abundance of within gene Shine-Dalgarno-like sequences had been shown to correlate negatively with protein production in *Methylobacterium extorquens* (Agashe et al. 2013) and we hypothesized that a high frequency of anti-SD-like sequences in our variants could reduce translation efficacy and thus resistance. No effect of SD-like sequences was found in *E. coli* and *P. aeruginosa* and contrary to expectations, we found a significant positive relationship between the abundance of anti-SD-like sequences and resistance in *A. baylyi*. This significance persisted even when the non-essential 5’ end of the coding sequence was removed, ruling out the potential positive effect of translation of a smaller, possibly more efficient protein. The biological determinant of this result is not clear and requires further examination. While the anti-SD sequence is commonly accepted to be conserved across bacterial species similarly to the conservation of the 16S rRNA sequence (Nakagawa et al. 2010), recent work (reviewed in Wen et al. 2021) puts into question the universality of the anti-SD sequence and the species-independent potential effect of within-gene anti-SD sequences by documenting within and between species anti-SD sequence diversity. We can thus not rule out that the anti-SD consensus sequence we used is not fully appropriate for *A. baylyi* or *P. aeruginosa*.

Our multi-species approach produced results suggesting that different species have different sensitivity to the various ways by which synonymous variation affects the phenotype. Whilst our results do not conclude that one codon index or sequence motif explains all of the phenotype diversity provided by synonymous variation, our findings do suggest that synonymous variation is acting on numerous levels simultaneously, in line with the diversity of synonymous mutation related effects published in recent years. Indeed, variant sequence composition acts on translational initiation, translation elongation, mRNA decay, protein folding and protein error rate (Kudla et al. 2009; Amorós-Moya et al. 2010; Cambray et al. 2018; Mittal et al. 2018; Walsh et al. 2020). Additionally, as transcription and translation are coupled in bacteria (Komar et al. 1999), translation-related positive or negative feedback on transcription might add another layer of complexity.

We reject the hypothesis that differences in synonymous variant phenotype are most explained by mRNA folding and its effects on translation initiation. Conservation of the first 30bps of the coding sequence did not prevent large differences in resistance phenotype and mRNA folding in the first 90 bps of the coding sequence did not explain significantly the resistance levels. We do not doubt the modulatory effect of this area and its impact on phenotype put forward in many recent papers (Kudla et al. 2009; Cambray et al. 2018; Shaferman et al. 2023). We do, however, note that it is only one of many synonymous sequence related effects on phenotype.

In conclusion, our multi-species approach revealed that synonymous variation affects the immediate success of HGT and is likely to be a factor orienting transfers. But our experimental results indicate that the compatibility between a variant and a species is mediated through a diversity of mechanisms, some of which not clearly understood yet, such that it is not possible for now to make direct prediction of resistance/activity level conferred from the nucleotide sequence of a gene. A full understanding of how the sequence characteristics determine the phenotype would allow for these sequence characteristics to be used as an input variable in general microbiota models (Coyte et al. 2022), to predict resistance gene and plasmid circulation. This study also pointed at species-specific plasmid copy number as a factor determining resistance level, immediate HGT success and the orientation of gene transfers.

## Materials and Methods

### Synonymous Variant Synthesis

Synonymous variants of the *aacC1* gene (GenBank: AAB20441.1, *Serratia marcescens*) were designed using the *Optimization Analysis* function of the COUSIN (COdon Usage Similarity INdex) tool (Bourret et al. 2019) and codon usage tables (CUT) of *E. coli* K12 MG1655 (GenBank: U00096.3), *P. aeruginosa* PAO1 (ATCC 15692, NC_002516.2), *A. baylyi* ADP1 (NC_005966.1) *and B. cereus* ATCC 14579 (GenBank: AP007209.1)*. B. cereus* had originally been chosen to include a Gram-positive species. However, the species was discontinued post-cloning after failure to express and maintain the pBBR1 plasmid and *aacC1* gene construct.

In each variant, the first 30 bps of the CDS were conserved to minimize the effects of differences in translation initiation around the translation start site linked to mRNA folding and ribosome access to the RBS. 20 variants were produced for each species using the random guided optimization option in which each synonymous codon is independently assigned based on the frequencies of use in the CUT (Tables S12-S16). An additional five variants were produced in *E. coli* using the reverse frequencies option, which reverses the frequencies of the input CUT. Four variants were produced using the one amino - one codon option, with each codon being coded by the most frequently used synonymous codon for each amino acid. This option was used once for each of the four species. An additional variant was produced in *E. coli* using the least frequent codon for each amino acid.

A subset of 31 variants was conserved from the 90 designed variants. Variants were chosen to capture the widest range of COUSIN values in all four species. The six variants with the highest COUSIN values in *A. baylyi* and *B. cereus* were chosen (Table S17). The six variants with the highest values were also chosen in *P. aeruginosa*. However, one variant was excluded due to high GC content related synthesis issues. The five highest values in *E. coli* were retained in addition to four variants produced using the reverse frequencies option. The five one amino acid - one codon option variants were also conserved. The reverse frequencies option was used to provide a better range of mismatch to codon usage in *E. coli* (Figure 1, S1). For full sequence information on the 31 variants and the wildtype, see tables S18-S22. Full gene sequences for these 31 variants and the wt variant were synthesized at TwistBioscience (San Francisco, California).

### pBBR1 plasmid

The pBBR1 plasmid was ordered from addgene (pBBR1-MCS-2 plasmid #85168). This broad host range plasmid contains a natural plasmid backbone consisting of a pBBR1 *OriV* and pBBR1 *Rep*. The plasmid also contains a kanamycin resistance gene and a multiple cloning site (MSC) (Kovach et al. 1995).

### Variant Cloning

The 32 synthesized gene fragments were cloned into the pBBR1 broad host range plasmid using restriction enzyme cloning techniques. The resulting plasmid were transformed into DH5a competent cells and selected on kanamycin 50µg/mL plates. Successful clones were verified by Sanger sequencing.

### Bacterial Transformation

Plasmids harboring *aacC1* synonymous variants were transformed into *A. baylyi (ADP1)*, *E. coli* (K12 MG1655) and *P. aeruginosa (PA01)* and plated on selective media. Bacterial clones were isolated, cultured in selective media and archived in glycerol at -70°C. Species specific transformation protocols are described in the supplementary methods.

### Bacterial growth curves

Samples were pre-cultured from frozen stocks for 24 hrs in LB without antibiotics. Samples were then transferred to a 96 well microplate containing serially diluted concentrations of gentamicin and a LB control. Growth curves were produced by incubating samples at 30°C (*A. baylyi*) or 37°C (*P. aeruginosa, E. coli*) for 24 hrs (48 hrs for *A. baylyi*) in a Spark Tecan spectrophotometer. A humidity cassette was employed to prevent evaporation. Growth was recorded by measuring absorbance at 600nm at T0 and approximately every 30 minutes for 24 hrs or 48 hrs. During each 30 minutes cycle, an absorbance measure was recorded, followed by a 22 minutes wait period, followed by a 5 minutes orbital shaking period (108 rpm, 2.5 mm amplitude). This was then followed by a 2 minutes settlement period. Growth curves were performed in triplicate.

### Growth curve analysis

Growth curve data was analyzed using the R package gcplyr (Blazanin 2023). The growth metrics AUC, carrying capacity (Max OD) and lag time were calculated from individual growth curves for each replicate, for each gentamicin concentration, for each variant and for each species. Individual growth metrics are available in Tables S23 to S25.

### Resistance levels (IC50 and MIC calculations)

IC50 (IC50 AUC) is defined here as the concentration of gentamicin that resulted in a 50% reduction in AUC compared to growth in the absence of antibiotics. IC50 was calculated using a four parameter log-logistic fit of the drm() function in R (drc package) MIC was evaluated from growth curve measures. We defined MIC as the lowest gentamicin concentration at which final OD was <0.1.

### Plasmid Copy Number

Plasmid copy number was measured by qPCR quantification of a 100 bp sequence of the 5’UTR of the *aacC1* gene and of the *RpoD* gene. *RpoD* is present in a single copy on the chromosome of each species. PCN was calculated with the following formula:

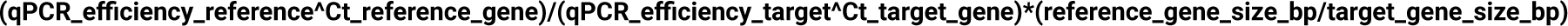

qPCR were conducted in triplicate on three *aacC1* variants (V2, V11, V19) in each of the three species. The qPCR conditions were as follows: LightCycler® 480 SYBR Green I Master (Roche), 45 cycles on LC480 (Roche) - 94°C for 10 seconds, followed by 66°C (rpoD primers) or 59°C (pBBR1 aacC1 primers) for 10 seconds, followed by 72°C for 10 seconds.

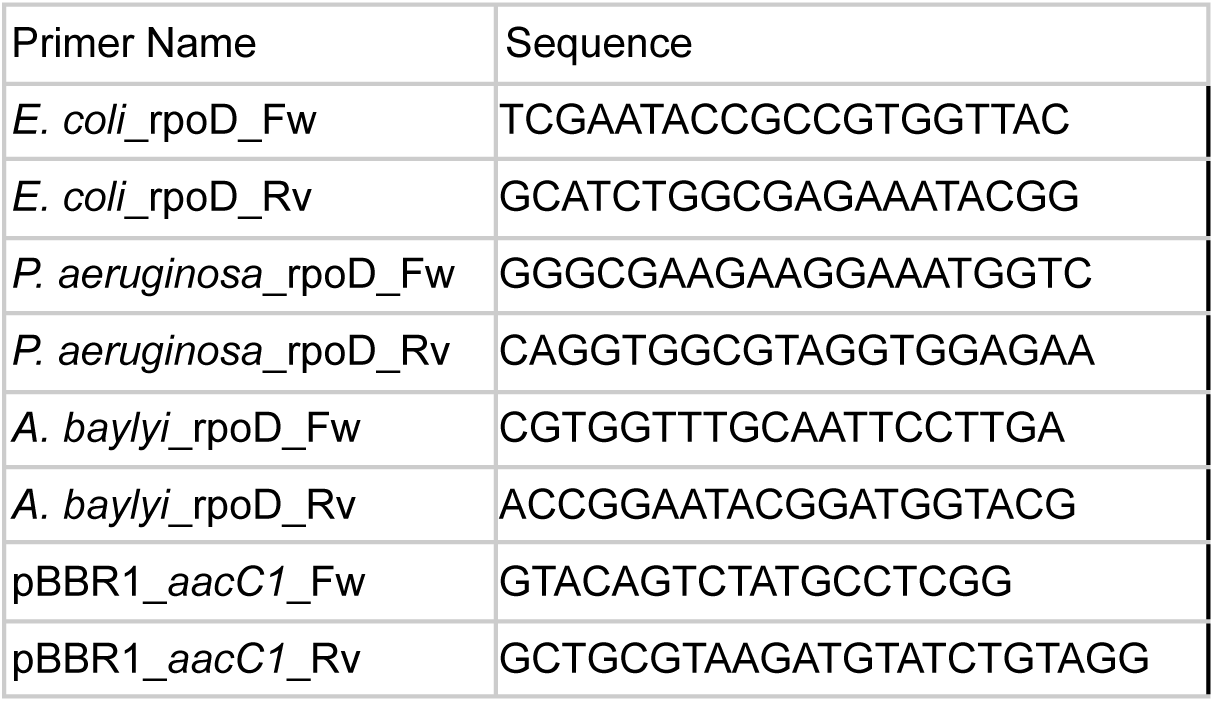

### Codon Usage Preferences Metric Calculations

COUSIN and CAI were calculated using the COUSIN tool (https://cousin.ird.fr/calculation.php), by inputting the CUT for each of the three species and the 32 variant coding sequences. CAI here is calculated on the frequencies of synonymous codons across the genome and not in a subset of highly expressed genes. COUSIN was calculated as described in Bourret et al. (2019).

tAI was calculated using the tai R package (https://github.com/mariodosreis/tai, <u>{Updating}</u>), by inputting tRNA copy number for each species and the variant coding sequences.

|Δ(GC3)| was calculated as the absolute value of the difference in GC3 between a variant sequence and the host genome. GC3 for each variant was obtained using the COUSIN tool calculation. GC3 for each species was taken from CoCoPuts described in Athey et al. (2017).

### Local codon usage metrics

The *aacC1* gene was split into 30 bp sliding windows (steps every 15 bps). CAI, COUSIN, tAI and |Δ(GC3)| were calculated for these 36 windows as above.

Protein structural domains of the *aacC1* gene coding sequence were extracted from https://www.ebi.ac.uk/pdbe/entry/pdb/1bo4/analysis (EMBL-EBI Protein Data Bank in Europe, 1bo4, (Wolf et al. 1998).

Bottleneck position and strength were calculated similarly to Navon and Pilpel (2011). Bottleneck position is defined here as the position of the window with the lowest tAI value. Bottleneck strength is defined here as the tAI of the bottleneck relative to the mean tAI of the 36 sliding windows (bottleneck tAI / mean tAI).

Rare codon chains were defined as the number of chains of 1, 2, 3, 4 or 5 rare codons, calculated using the str_count() in R. We define rare codons for each species as codons that are underrepresented in the host CUT. More specifically, we defined rare codons as those with less than half their expected frequency under the null hypothesis of equal frequency within a synonymous codon family (ex. for a codon within a four synonymous codon families, a rare codon is a codon with a frequency <0.125).

### Shine-Dalgarno-like sequences

Shine-Dalgarno-like sequences were defined as hexamers with high affinity to the conserved anti-SD sequence. This was done by calculating the free binding energy between CACCUCCU and all possible hexamers within the *aacC1* gene variants using the RNAup function of the ViennaRNA 2.0 Package (Lorenz et al. 2011). We counted the number of hexamers in each synonymous variant with a binding free energy < -4 kcal/mol which is the definition of a weak SD-like in (Hockenberry et al. 2018). A strong SD-like sequence is defined as < -7 kcal/mol.

### mRNA secondary structure

mRNA folding energy was calculated using the Vienna RNA 2.0 function RNAfold, by inputting the -30:60 bp section of each variant mRNA sequence (1:3 = ATG of the coding sequence).

### Statistical Tests

All one-way and two-way ANOVA tests and Pearson Correlation tests were performed in R using the functions aov() and cor.test() (stats package), respectively.

*P*-values were corrected using the Benjamini-Hochberg (BH) procedure using the p.adjust() function (stats package) in R.

*The data underlying this article are available in the article and in its online supplementary material*.

## Acknowledgements

We thank Daniel Weinreich, Sylvain Gandon, Guillaume Cambray and Benoît Nabholz for stimulating scientific and technical discussions. This work was supported by the ERC HGTCODONUSE (ERC-2015-CoG-682819) to SB.

## Bibliography

1. Acar Kirit, Hande, Mato Lagator, and Jonathan P. Bollback. 2020. “Experimental Determination of Evolutionary Barriers to Horizontal Gene Transfer.” BMC Microbiology 20 (1): 326. 10.1186/s12866-020-01983-5.

2. Agashe, Deepa, N. Cecilia Martinez-Gomez, D. Allan Drummond, and Christopher J. Marx. 2013. “Good Codons, Bad Transcript: Large Reductions in Gene Expression and Fitness Arising from Synonymous Mutations in a Key Enzyme.” Molecular Biology and Evolution 30 (3): 549–60. 10.1093/molbev/mss273.

3. Agashe, Deepa, Mrudula Sane, Kruttika Phalnikar, Gaurav D. Diwan, Alefiyah Habibullah, Norma Cecilia Martinez-Gomez, Vinaya Sahasrabuddhe, et al. 2016. “Large-Effect Beneficial Synonymous Mutations Mediate Rapid and Parallel Adaptation in a Bacterium.” Molecular Biology and Evolution 33 (6): 1542–53. 10.1093/molbev/msw035.

4. Aguirre, Beatriz, Miguel Costas, Nallely Cabrera, Guillermo Mendoza-Hernández, Donald L. Helseth, Paulette Fernández, Marietta Tuena de Gómez-Puyou, Ruy Pérez-Montfort, Alfredo Torres-Larios, and Armando Gómez Puyou. 2011. “A Ribosomal Misincorporation of Lys for Arg in Human Triosephosphate Isomerase Expressed in Escherichia Coli Gives Rise to Two Protein Populations.” PLoS ONE 6 (6). 10.1371/journal.pone.0021035.

5. Alonso-del Valle, Aida, Laura Toribio-Celestino, Anna Quirant, Carles Tardio Pi, Javier DelaFuente, Rafael Canton, Eduardo P. C. Rocha, Carles Ubeda, Rafael Peña-Miller, and Alvaro San Millan. 2023. “Antimicrobial Resistance Level and Conjugation Permissiveness Shape Plasmid Distribution in Clinical Enterobacteria.” Proceedings of the National Academy of Sciences 120 (51): e2314135120. 10.1073/pnas.2314135120.

6. Amorós-Moya, Dolors, Stéphanie Bedhomme, Marita Hermann, and Ignacio G. Bravo. 2010. “Evolution in Regulatory Regions Rapidly Compensates the Cost of Nonoptimal Codon Usage.” Molecular Biology and Evolution 27 (9): 2141–51. 10.1093/molbev/msq103.

7. Arnold, Brian J., I.-Ting Huang, and William P. Hanage. 2022. “Horizontal Gene Transfer and Adaptive Evolution in Bacteria.” Nature Reviews Microbiology 20 (4): 206–18. 10.1038/s41579-021-00650-4.

8. Athey, John, Aikaterini Alexaki, Ekaterina Osipova, Alexandre Rostovtsev, Luis V. Santana-Quintero, Upendra Katneni, Vahan Simonyan, and Chava Kimchi-Sarfaty. 2017. “A New and Updated Resource for Codon Usage Tables.” BMC Bioinformatics 18 (September): 391. 10.1186/s12859-017-1793-7.

9. Bahiri-Elitzur, Shir, and Tamir Tuller. 2021. “Codon-Based Indices for Modeling Gene Expression and Transcript Evolution.” Computational and Structural Biotechnology Journal 19 (January): 2646–63. 10.1016/j.csbj.2021.04.042.

10. Bailey, Susan F, Luz Angela Alonso Morales, and Rees Kassen. 2021. “Effects of Synonymous Mutations beyond Codon Bias: The Evidence for Adaptive Synonymous Substitutions from Microbial Evolution Experiments.” Genome Biology and Evolution 13 (9): evab141. 10.1093/gbe/evab141.

11. Bailey, Susan F., Aaron Hinz, and Rees Kassen. 2014. “Adaptive Synonymous Mutations in an Experimentally Evolved Pseudomonas Fluorescens Population.” Nature Communications 5 (1): 4076. 10.1038/ncomms5076.

12. Baltrus, David A. 2013. “Exploring the Costs of Horizontal Gene Transfer.” Trends in Ecology and Evolution 28 (8): 489–95. 10.1016/j.tree.2013.04.002.

13. Blazanin, Michael. 2023. “Gcplyr: An R Package for Microbial Growth Curve Data Analysis.” bioRxiv. 10.1101/2023.04.30.538883.

14. Bourret, Jérôme, Samuel Alizon, and Ignacio G. Bravo. 2019. “COUSIN (COdon Usage Similarity INdex): A Normalized Measure of Codon Usage Preferences.” Genome Biology and Evolution 11 (12): 3523–28. 10.1093/gbe/evz262.

15. Brito, Ilana Lauren. 2021. “Examining Horizontal Gene Transfer in Microbial Communities.” *Nature Reviews Microbiology*, April. 10.1038/s41579-021-00534-7.

16. Buchan, J. Ross, Lorna S. Aucott, and Ian Stansfield. 2006. “tRNA Properties Help Shape Codon Pair Preferences in Open Reading Frames.” Nucleic Acids Research 34 (3): 1015–27. 10.1093/nar/gkj488.

17. Burch, Christina L., Artur Romanchuk, Michael Kelly, Yingfang Wu, and Corbin D. Jones. 2022. “Genome-Wide Determination of Barriers to Horizontal Gene Transfer.” bioRxiv. 10.1101/2022.06.29.498157.

18. Callens, Martijn, Léa Pradier, Michael Finnegan, Caroline Rose, and Stéphanie Bedhomme. n.d. “Title: Read between the Lines: Diversity of Non-Translational Selection Pressures on Local Codon Usage.” 10.1093/gbe/evab097/6263832.

19. Callens, Martijn, Celine Scornavacca, and Stéphanie Bedhomme. 2021. “Evolutionary Responses to Codon Usage of Horizontally Transferred Genes in Pseudomonas Aeruginosa: Gene Retention, Amelioration and Compensatory Evolution.” Microbial Genomics 7 (6): 000587. 10.1099/mgen.0.000587.

20. Cambray, Guillaume, Joao C. Guimaraes, and Adam Paul Arkin. 2018. “Evaluation of 244,000 Synthetic Sequences Reveals Design Principles to Optimize Translation in Escherichia Coli.” Nature Biotechnology 36 (10): 1005–1005. 10.1038/nbt.4238.

21. Chiang, C. S., Y. C. Xu, and H. Bremer. 1991. “Role of DnaA Protein during Replication of Plasmid pBR322 in Escherichia Coli.” Molecular & General Genetics: MGG 225 (3): 435–42. 10.1007/BF00261684.

22. Chu, Hoi Yee, Kathleen Sprouffske, and Andreas Wagner. 2018. “Assessing the Benefits of Horizontal Gene Transfer by Laboratory Evolution and Genome Sequencing.” BMC Evolutionary Biology 18 (April): 54. 10.1186/s12862-018-1164-7.

23. Coyte, Katharine Z., Cagla Stevenson, Christopher G. Knight, Ellie Harrison, James P. J. Hall, and Michael A. Brockhurst. 2022. “Horizontal Gene Transfer and Ecological Interactions Jointly Control Microbiome Stability.” PLoS Biology 20 (11): e3001847. 10.1371/journal.pbio.3001847.

24. Del Solar, Gloria, and Manuel Espinosa. 2000. “Plasmid Copy Number Control: An Ever-Growing Story.” Molecular Microbiology 37 (3): 492–500. 10.1046/j.1365-2958.2000.02005.x.

25. Didelot, Xavier, Sandra Nell, Ines Yang, Sabrina Woltemate, Schalk van der Merwe, and Sebastian Suerbaum. 2013. “Genomic Evolution and Transmission of Helicobacter Pylori in Two South African Families.” Proceedings of the National Academy of Sciences of the United States of America 110 (34): 13880–85. 10.1073/pnas.1304681110.

26. Drummond, D Allan, and Claus O Wilke. 2010. “NIH Public Access” 10(10): 715–24. 10.1038/nrg2662.The.

27. Ederth, J., Leif A. Isaksson, and F. Abdulkarim. 2002. “Origin-Specific Reduction of ColE1 Plasmid Copy Number Due to Mutations in a Distinct Region of the Escherichia Coli RNA Polymerase.” Molecular Genetics and Genomics: MGG 267 (5): 587–92. 10.1007/s00438-002-0689-y.

28. Forsberg, Kevin J., Sanket Patel, Molly K. Gibson, Christian L. Lauber, Rob Knight, Noah Fierer, and Gautam Dantas. 2014. “Bacterial Phylogeny Structures Soil Resistomes across Habitats.” Nature 509 (7502): 612–16. 10.1038/nature13377.

29. Fragata, Inês, Sebastian Matuszewski, Mark A. Schmitz, Thomas Bataillon, Jeffrey D. Jensen, and Claudia Bank. 2018. “The Fitness Landscape of the Codon Space across Environments.” Heredity 121 (5): 422–37. 10.1038/s41437-018-0125-7.

30. Frazão, Nelson, Ana Sousa, Michael Lässig, and Isabel Gordo. 2019. “Horizontal Gene Transfer Overrides Mutation in Escherichia Coli Colonizing the Mammalian Gut.” Proceedings of the National Academy of Sciences 116 (36): 17906–15. 10.1073/pnas.1906958116.

31. Gomes, Antonio L. C., Nathan I. Johns, Anthony Yang, Florencia Velez-Cortes, Christopher S. Smillie, Mark B. Smith, Eric J. Alm, and Harris H. Wang. 2020. “Genome and Sequence Determinants Governing the Expression of Horizontally Acquired DNA in Bacteria.” The ISME Journal 14 (9): 2347–57. 10.1038/s41396-020-0696-1.

32. Granato, Elisa T., Thomas A. Meiller-Legrand, and Kevin R. Foster. 2019. “The Evolution and Ecology of Bacterial Warfare.” Current Biology 29 (11): R521–37. 10.1016/j.cub.2019.04.024.

33. Grantham, R, C Gautier, M Gouy, R Mercier, and A Pavé. 1980. “Codon Catalog Usage and the Genome Hypothesis.” Nucleic Acids Research 8 (1): r49–62. https://www.ncbi.nlm.nih.gov/pmc/articles/PMC327256/.

34. Harrison, Ellie, and Michael A. Brockhurst. 2012. “Plasmid-Mediated Horizontal Gene Transfer Is a Coevolutionary Process.” Trends in Microbiology 20 (6): 262–67. 10.1016/j.tim.2012.04.003.

35. Hershberg, Ruth, and Dmitri A. Petrov. 2008. “Selection on Codon Bias.” Annual Review of Genetics 42: 287–99. 10.1146/annurev.genet.42.110807.091442.

36. Hockenberry, Adam J., Michael C. Jewett, Luıs A.N. Amaral, and Claus O. Wilke. 2018. “Within-Gene Shine-Dalgarno Sequences Are Not Selected for Function.” Molecular Biology and Evolution 35 (10): 2487–98. 10.1093/molbev/msy150.

37. Horton, James S., Louise M. Flanagan, Robert W. Jackson, Nicholas K. Priest, and Tiffany B. Taylor. 2021. “A Mutational Hotspot That Determines Highly Repeatable Evolution Can Be Built and Broken by Silent Genetic Changes.” Nature Communications 12 (1): 6092. 10.1038/s41467-021-26286-9.

38. Huai, Wei, Qing-Bian Ma, Jia-Jia Zheng, Yang Zhao, and Qiang-Rong Zhai. 2019. “Distribution and Drug Resistance of Pathogenic Bacteria in Emergency Patients.” World Journal of Clinical Cases 7 (20): 3175–84. 10.12998/wjcc.v7.i20.3175.

39. Komar, A. A., T. Lesnik, and C. Reiss. 1999. “Synonymous Codon Substitutions Affect Ribosome Traffic and Protein Folding during in Vitro Translation.” FEBS Letters 462 (3): 387–91. 10.1016/s0014-5793(99)01566-5.

40. Kovach, M. E., P. H. Elzer, D. S. Hill, G. T. Robertson, M. A. Farris, R. M. Roop, and K. M. Peterson. 1995. “Four New Derivatives of the Broad-Host-Range Cloning Vector pBBR1MCS, Carrying Different Antibiotic-Resistance Cassettes.” Gene 166 (1): 175–76. 10.1016/0378-1119(95)00584-1.

41. Kowalska-Krochmal, Beata, and Ruth Dudek-Wicher. 2021. “The Minimum Inhibitory Concentration of Antibiotics: Methods, Interpretation, Clinical Relevance.” Pathogens 10 (2): 165. 10.3390/pathogens10020165.

42. Kudla, Grzegorz, Andrew W. Murray, David Tollervey, and Joshua B. Plotkin. 2009. “Coding-Sequence Determinants of Expression in Escherichia Coli.” Science 324 (5924): 255–58. 10.1126/science.1170160.

43. Lebeuf-Taylor, Eleonore, Nick McCloskey, Susan F. Bailey, Aaron Hinz, and Rees Kassen. 2019. “The Distribution of Fitness Effects among Synonymous Mutations in a Gene under Directional Selection.” eLife 8 (July): e45952. 10.7554/eLife.45952.

44. Lind, Peter A., Christina Tobin, Otto G. Berg, Charles G. Kurland, and Dan I. Andersson. 2010. “Compensatory Gene Amplification Restores Fitness after Inter-Species Gene Replacements.” Molecular Microbiology 75 (5): 1078–89. 10.1111/j.1365-2958.2009.07030.x.

45. Liu, Yi, Qian Yang, and Fangzhou Zhao. 2021. “Synonymous but Not Silent: The Codon Usage Code for Gene Expression and Protein Folding.” Annual Review of Biochemistry 90 (June): 375–401. 10.1146/annurev-biochem-071320-112701.

46. Lopilato, J., S. Bortner, and J. Beckwith. 1986. “Mutations in a New Chromosomal Gene of Escherichia Coli K-12, pcnB, Reduce Plasmid Copy Number of pBR322 and Its Derivatives.” Molecular & General Genetics: MGG 205 (2): 285–90. 10.1007/BF00430440.

47. Lorenz, Ronny, Stephan H. Bernhart, Christian Höner zu Siederdissen, Hakim Tafer, Christoph Flamm, Peter F. Stadler, and Ivo L. Hofacker. 2011. “ViennaRNA Package 2.0.” Algorithms for Molecular Biology 6 (1): 26. 10.1186/1748-7188-6-26.

48. Martens, Andrew T., James Taylor, and Vincent J. Hilser. 2015. “Ribosome A and P Sites Revealed by Length Analysis of Ribosome Profiling Data.” Nucleic Acids Research 43 (7): 3680–87. 10.1093/nar/gkv200.

49. Mittal, Pragya, James Brindle, Julie Stephen, Joshua B. Plotkin, and Grzegorz Kudla. 2018. “Codon Usage Influences Fitness through RNA Toxicity.” Proceedings of the National Academy of Sciences of the United States of America 115 (34): 8639–44. 10.1073/pnas.1810022115.

50. Mordret, Ernest, Orna Dahan, Omer Asraf, Roni Rak, Avia Yehonadav, Georgina D. Barnabas, Jürgen Cox, T. Geiger, Ariel B. Lindner, and Yitzhak Pilpel. 2019. “Systematic Detection of Amino Acid Substitutions in Proteomes Reveals Mechanistic Basis of Ribosome Errors and Selection for Translation Fidelity.” Molecular Cell 75 (3): 427–441.e5. 10.1016/j.molcel.2019.06.041.

51. Murray, Christopher J. L., Kevin Shunji Ikuta, Fablina Sharara, Lucien Swetschinski, Gisela Robles Aguilar, Authia Gray, Chieh Han, et al. 2022. “Global Burden of Bacterial Antimicrobial Resistance in 2019: A Systematic Analysis.” The Lancet 399 (10325): 629–55. 10.1016/S0140-6736(21)02724-0.

52. Nakagawa, So, Yoshihito Niimura, Kin-ichiro Miura, and Takashi Gojobori. 2010. “Dynamic Evolution of Translation Initiation Mechanisms in Prokaryotes.” Proceedings of the National Academy of Sciences 107 (14): 6382–87. 10.1073/pnas.1002036107.

53. Navon, Sivan, and Yitzhak Pilpel. 2011. “The Role of Codon Selection in Regulation of Translation Efficiency Deduced from Synthetic Libraries.” Genome Biology 12 (2): R12. 10.1186/gb-2011-12-2-r12.

54. Nordström, Kurt. 2006. “Plasmid R1—Replication and Its Control.” Plasmid 55 (1): 1–26. 10.1016/j.plasmid.2005.07.002.

55. Pena-Gonzalez, Angela, Luis M. Rodriguez-R, Chung K. Marston, Jay E. Gee, Christopher A. Gulvik, Cari B. Kolton, Elke Saile, Michael Frace, Alex R. Hoffmaster, and Konstantinos T. Konstantinidis. 2018. “Genomic Characterization and Copy Number Variation of Bacillus Anthracis Plasmids pXO1 and pXO2 in a Historical Collection of 412 Strains.” mSystems 3 (4): 10.1128/msystems.00065-18.

56. Podolsky, Scott H. 2018. “The Evolving Response to Antibiotic Resistance (1945–2018).” Palgrave Communications 4 (1): 1–8. 10.1057/s41599-018-0181-x.

57. Popa, Ovidiu, and Tal Dagan. 2011. “Trends and Barriers to Lateral Gene Transfer in Prokaryotes.” *Current Opinion in Microbiology*, Antimicrobials/Genomics, 14 (5): 615–23. 10.1016/j.mib.2011.07.027.

58. Pradier, Léa, and Stéphanie Bedhomme. 2023. “Ecology, More than Antibiotics Consumption, Is the Major Predictor for the Global Distribution of Aminoglycoside-Modifying Enzymes.” Edited by Gabriel Perron and Meredith C Schuman. eLife 12 (February): e77015. 10.7554/eLife.77015.

59. Ram, Yoav, Eynat Dellus-Gur, Maayan Bibi, Kedar Karkare, Uri Obolski, Marcus W. Feldman, Tim F. Cooper, Judith Berman, and Lilach Hadany. 2019. “Predicting Microbial Growth in a Mixed Culture from Growth Curve Data.” Proceedings of the National Academy of Sciences 116 (29): 14698–707. 10.1073/pnas.1902217116.

60. Reis, Mario dos, Lorenz Wernisch, and Renos Savva. 2003. “Unexpected Correlations between Gene Expression and Codon Usage Bias from Microarray Data for the Whole Escherichia Coli K-12 Genome.” Nucleic Acids Research 31 (23): 6976–85. 10.1093/nar/gkg897.

61. Rocha, Eduardo P.C. 2004. “Codon Usage Bias from tRNA’s Point of View: Redundancy, Specialization, and Efficient Decoding for Translation Optimization.” Genome Research 14 (11): 2279–86. 10.1101/gr.2896904.

62. Rodriguez-Beltran, Jeronimo, J. Carlos R. Hernandez-Beltran, Javier Delafuente, Jose A. Escudero, Ayari Fuentes-Hernandez, R. Craig MacLean, Rafael Peña-Miller, and Alvaro San Millan. 2018. “Multicopy Plasmids Allow Bacteria to Escape from Fitness Trade-Offs during Evolutionary Innovation.” Nature Ecology and Evolution 2 (5): 873–81. 10.1038/s41559-018-0529-z.

63. San Millan, Alvaro, Macarena Toll-Riera, Qin Qi, and R. Craig MacLean. 2015. “Interactions between Horizontally Acquired Genes Create a Fitness Cost in Pseudomonas Aeruginosa.” Nature Communications 6 (1): 6845. 10.1038/ncomms7845.

64. Santucci, R. A., and J. N. Krieger. 2000. “Gentamicin for the Practicing Urologist: Review of Efficacy, Single Daily Dosing and ‘Switch’ Therapy.” The Journal of Urology 163 (4): 1076–84. 10.1016/s0022-5347(05)67697-5.

65. Schmitz, Alexander, and Fuzhong Zhang. 2021. “Massively Parallel Gene Expression Variation Measurement of a Synonymous Codon Library.” BMC Genomics 22 (1): 149. 10.1186/s12864-021-07462-z.

66. Shaferman, Michael, Melis Gencel, Noga Alon, Khawla Alasad, Barak Rotblat, Adrian W R Serohijos, Lital Alfonta, and Shimon Bershtein. 2023. “The Fitness Effects of Codon Composition of the Horizontally Transferred Antibiotic Resistance Genes Intensify at Sub-Lethal Antibiotic Levels.” Molecular Biology and Evolution 40 (6): msad123. 10.1093/molbev/msad123.

67. Sharpl, Paul M, and Wen-Hsiung Li2. n.d. “Nucleic Acids Research The Codon Adaptation Index-a Measure of Directional Synonymous Codon Usage Bias, and Its Potential Applications.” Vol. 15.

68. Stoletzki, Nina, and Adam Eyre-Walker. 2007. “Synonymous Codon Usage in Escherichia Coli: Selection for Translational Accuracy.” Molecular Biology and Evolution 24 (2): 374–81. 10.1093/molbev/msl166.

69. Thomas, Christopher M., and Kaare M. Nielsen. 2005. “Mechanisms of, and Barriers to, Horizontal Gene Transfer between Bacteria.” Nature Reviews Microbiology 3 (9): 711–21. 10.1038/nrmicro1234.

70. Tian, Jian, Yaru Yan, Qingxia Yue, Xiaoqing Liu, Xiaoyu Chu, Ningfeng Wu, and Yunliu Fan. 2017. “Predicting Synonymous Codon Usage and Optimizing the Heterologous Gene for Expression in E. Coli.” Scientific Reports 7 (1): 9926. 10.1038/s41598-017-10546-0.

71. Verma, Manasvi, Junhong Choi, Kyle A. Cottrell, Zeno Lavagnino, Erica N. Thomas, Slavica Pavlovic-Djuranovic, Pawel Szczesny, et al. 2019. “A Short Translational Ramp Determines the Efficiency of Protein Synthesis.” Nature Communications 10 (1): 5774. 10.1038/s41467-019-13810-1.

72. Walsh, Ian M, Micayla A Bowman, F Soto Santarriaga, Anabel Rodriguez, and Patricia L Clark. n.d. “Synonymous Codon Substitutions Perturb Cotranslational Protein Folding in Vivo and Impair Cell Fitness.” 10.1073/pnas.1907126117/-/DCSupplemental.

73. Wen, Jin-Der, Syue-Ting Kuo, and Hsin-Hung David Chou. 2021. “The Diversity of Shine-Dalgarno Sequences Sheds Light on the Evolution of Translation Initiation.” RNA Biology 18 (11): 1489–1500. 10.1080/15476286.2020.1861406.

74. Wolf, Eva, Alex Vassilev, Yasutaka Makino, Andrej Sali, Yoshihiro Nakatani, and Stephen K. Burley. 1998. “Crystal Structure of a GCN5-Related N-Acetyltransferase: Serratia Marcescens Aminoglycoside 3-N-Acetyltransferase.” Cell 94 (4): 439–49. 10.1016/S0092-8674(00)81585-8.

75. Woods, Laura C, Rebecca J Gorrell, Frank Taylor, Tim Connallon, Terry Kwok, and Michael J McDonald. n.d. “Horizontal Gene Transfer Potentiates Adaptation by Reducing Selective Constraints on the Spread of Genetic Variation.” 10.1073/pnas.2005331117/-/DCSupplemental.

76. Xu, F., S. Lin-Chao, and S. N. Cohen. 1993. “The Escherichia Coli pcnB Gene Promotes Adenylylation of Antisense RNAI of ColE1-Type Plasmids in Vivo and Degradation of RNAI Decay Intermediates.” Proceedings of the National Academy of Sciences of the United States of America 90 (14): 6756–60. 10.1073/pnas.90.14.6756.

77. Yan, Xiaowei, Tim A. Hoek, Ronald D. Vale, and Marvin E. Tanenbaum. 2016. “Dynamics of Translation of Single mRNA Molecules in Vivo.” Cell 165 (4): 976–89. 10.1016/j.cell.2016.04.034.

78. Yang, Y. L., and B. Polisky. 1999. “Allele-Specific Suppression of ColE1 High-Copy-Number Mutants by a rpoB Mutation of Escherichia Coli.” Plasmid 41 (1): 55–62. 10.1006/plas.1998.1378.

79. Yannai, Adi, Sophia Katz, and Ruth Hershberg. 2018. “The Codon Usage of Lowly Expressed Genes Is Subject to Natural Selection.” Genome Biology and Evolution 10 (5): 1237–46. 10.1093/gbe/evy084.

80. Zhu, Ying, Wei E Huang, and Qiwen Yang. 2022. “Clinical Perspective of Antimicrobial Resistance in Bacteria.” Infection and Drug Resistance 15 (March): 735–46. 10.2147/IDR.S345574.

