## Supplementary Figures S1-S8, Table S1 for "Hurdles to Horizontal Gene Transfer: Synonymous variation determines antibiotic resistance phenotype across species"

### Supplementary Materials

#### Supplementary Materials and Methods:

*E. coli* K12 MG1655 competent cells were mixed with 5ng of plasmid DNA for 15 minutes @4°C after which the samples were subject to 2000V for 5ms in an Eppendorf Eporator (VWR). Samples were recovered in 1 ml of LB media for 1hr @37°C, 220 rpm. Samples were then grown overnight on selective media (kanamycin 50 μg/mL). A single colony was isolated, grown overnight in 4 ml of LB and archived in glycerol at -80°C.

*P. aeruginosa* PAO1 (ATCC 15692) competent cells were mixed with 5ng plasmid DNA for 15 minutes @4°C after which the samples were subject to 2500V for 5ms in an Eppendorf Eporator (VWR). Samples were recovered in 1 ml of LB media for 1hr at 37°C, 220rpm. Samples were then grown overnight on selective media (gentamicin 50 μg/mL). A single colony was isolated, grown overnight in 4 ml of LB and archived in glycerol at -80°C.

*A. baylyi* ADP1 was cultured from frozen glycerol stocks overnight at 30°C, 220rpm. 1 ml of fresh culture was mixed with 100 ng of plasmid DNA and incubated overnight at 30°C, 220rpm, exploiting *A. baylyi*'s natural competency. The mix was then plated on selective plates (kanamycin 10μg/mL) and grown overnight at 30°C, 220rpm. A single colony was isolated, grown overnight in 4 ml of LB and archived in glycerol at -80°C.

#### Supplementary Figures

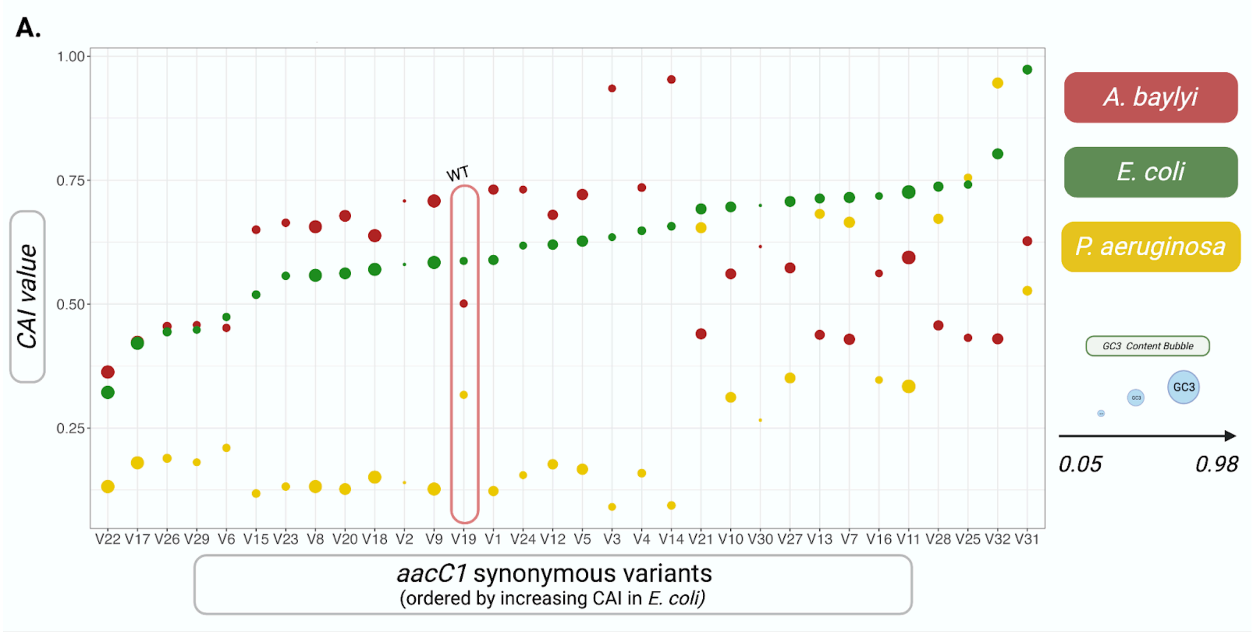


**Figure S1: Codon Adaptation Index (CAI) for the 32 synonymous *aacC1* variants.** Variants are ordered by increasing CAI values in *E. coli*. GC content at the third base of each codon (GC3) is represented by bubble size.

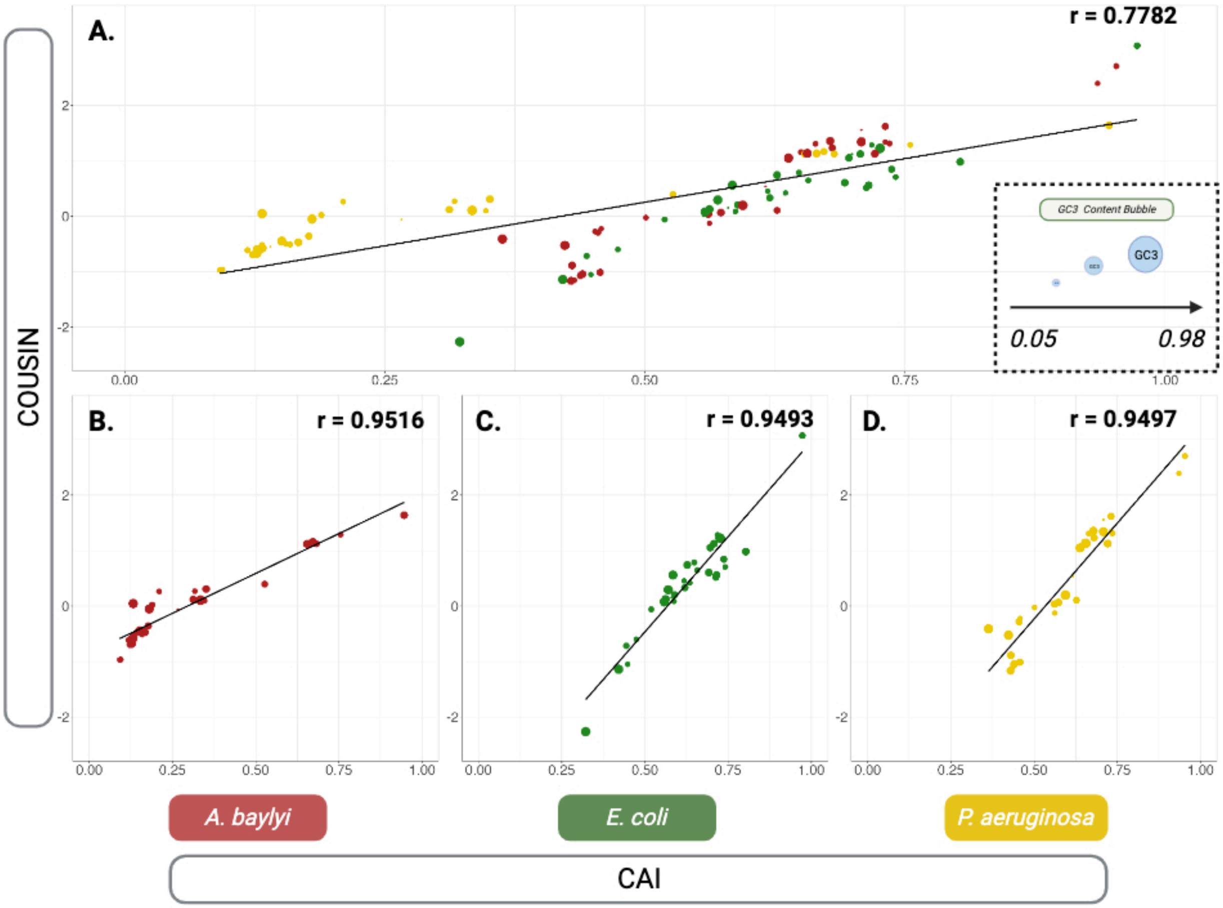


**Figure S2: Correlation between CAI and COUSIN for the 32 *aacC1* synonymous variants.** (A) Pearson correlation between CAI and COUSIN in all species. GC3 content is represented by bubble size. (B) Pearson correlation between CAI and COUSIN in all *A. baylyi*. (C) Pearson correlation between CAI and COUSIN in all *E. coli*. (D) Pearson correlation between CAI and COUSIN in all *P. aeruginosa*.

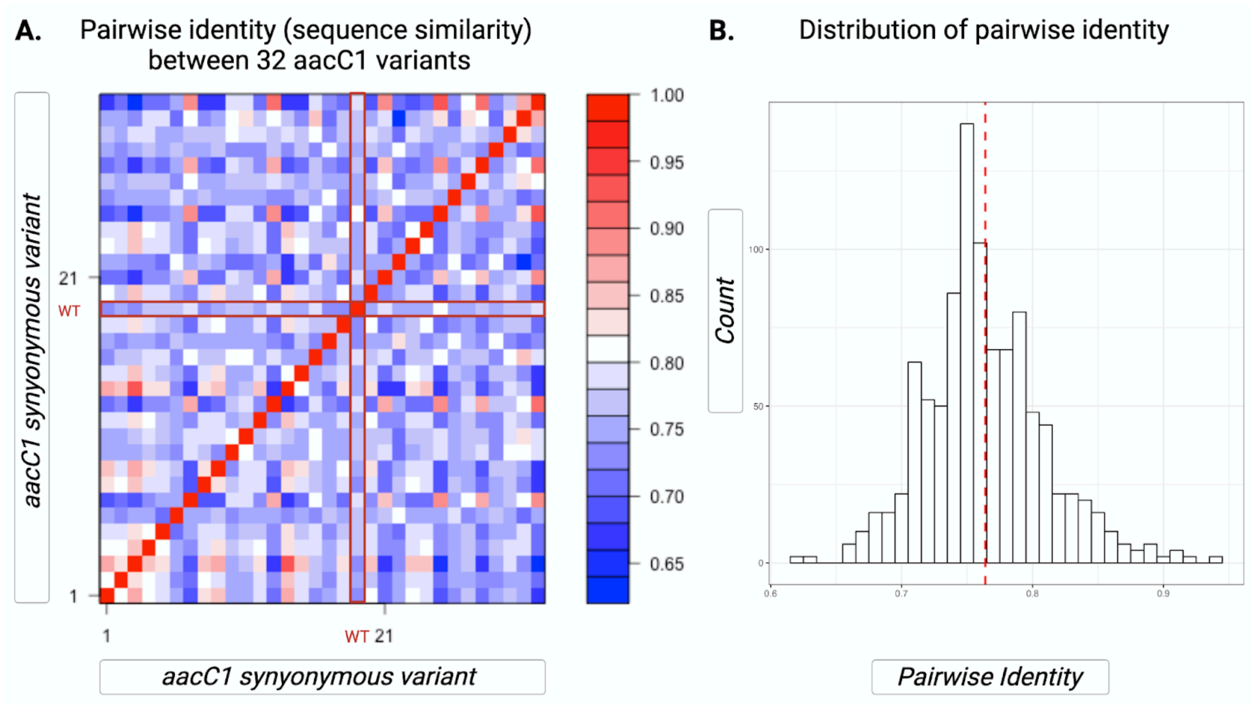


**Figure S3: Pairwise identity of the 32 *aacC1* synonymous variants:** (A) Heatmap of *aacC1* variants pairwise identity. Pairwise distance between the wt *aacC1* sequence, and the 31 synonymous variants is boxed in red. (B) Distribution of pairwise identities between the 32 synonymous variants. The red dotted vertical line represents the median pairwise distance.

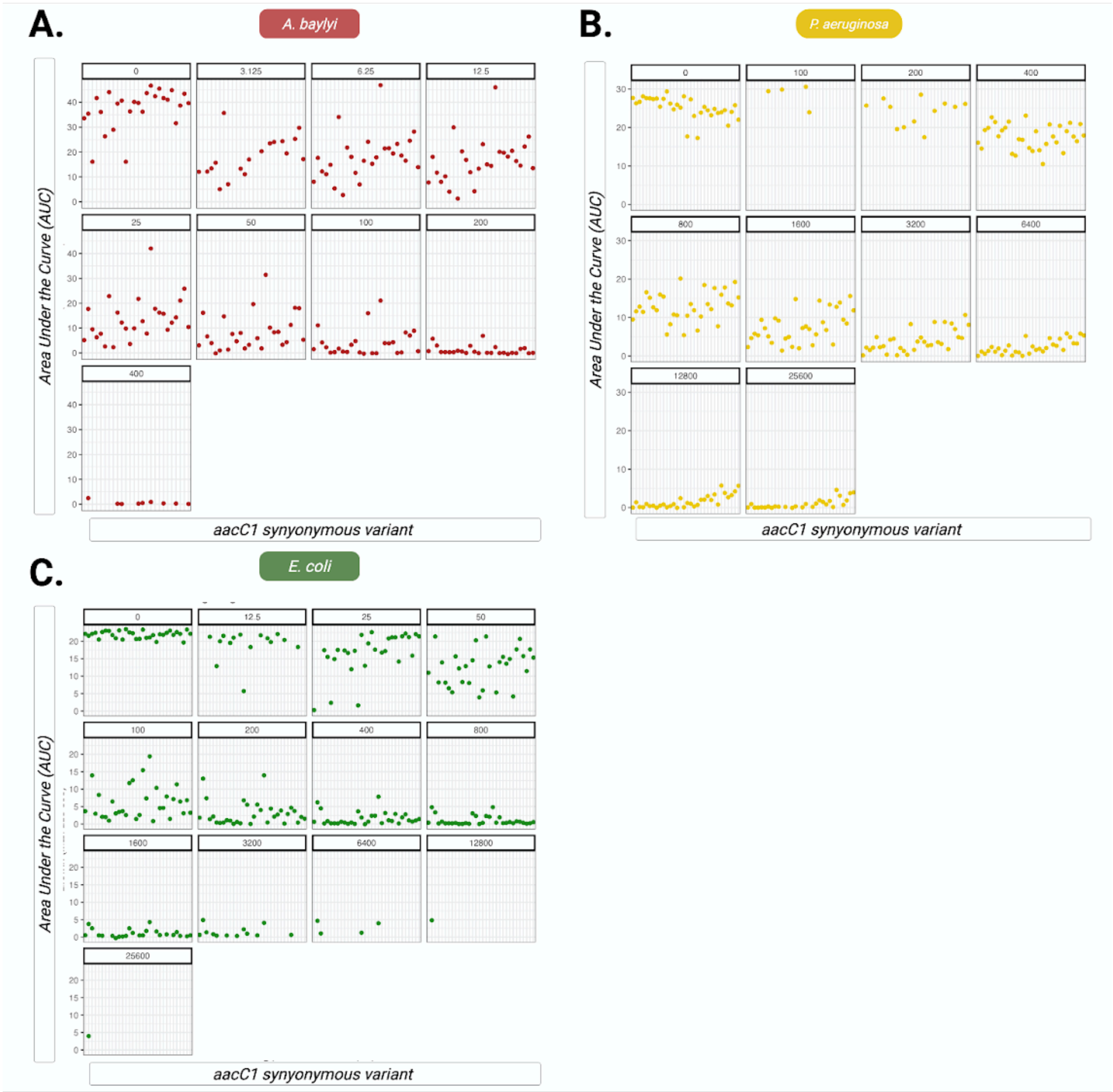


**Figure S4. Area Under the Curve (AUC):** Mean AUC values for each variant in each gentamicin concentration. Variants are ordered from 1-32 on the x avis. The wt variant is V19. **A.** *A. baylyi.* The graphs do not contain V3, V4, V12, V14, V20 or V32 as they could not be transformed into *A. baylyi*. **B.** *E. coli* **C.** *P. aeruginosa.* AUC of individual samples is available in supplementary file S24-S26.

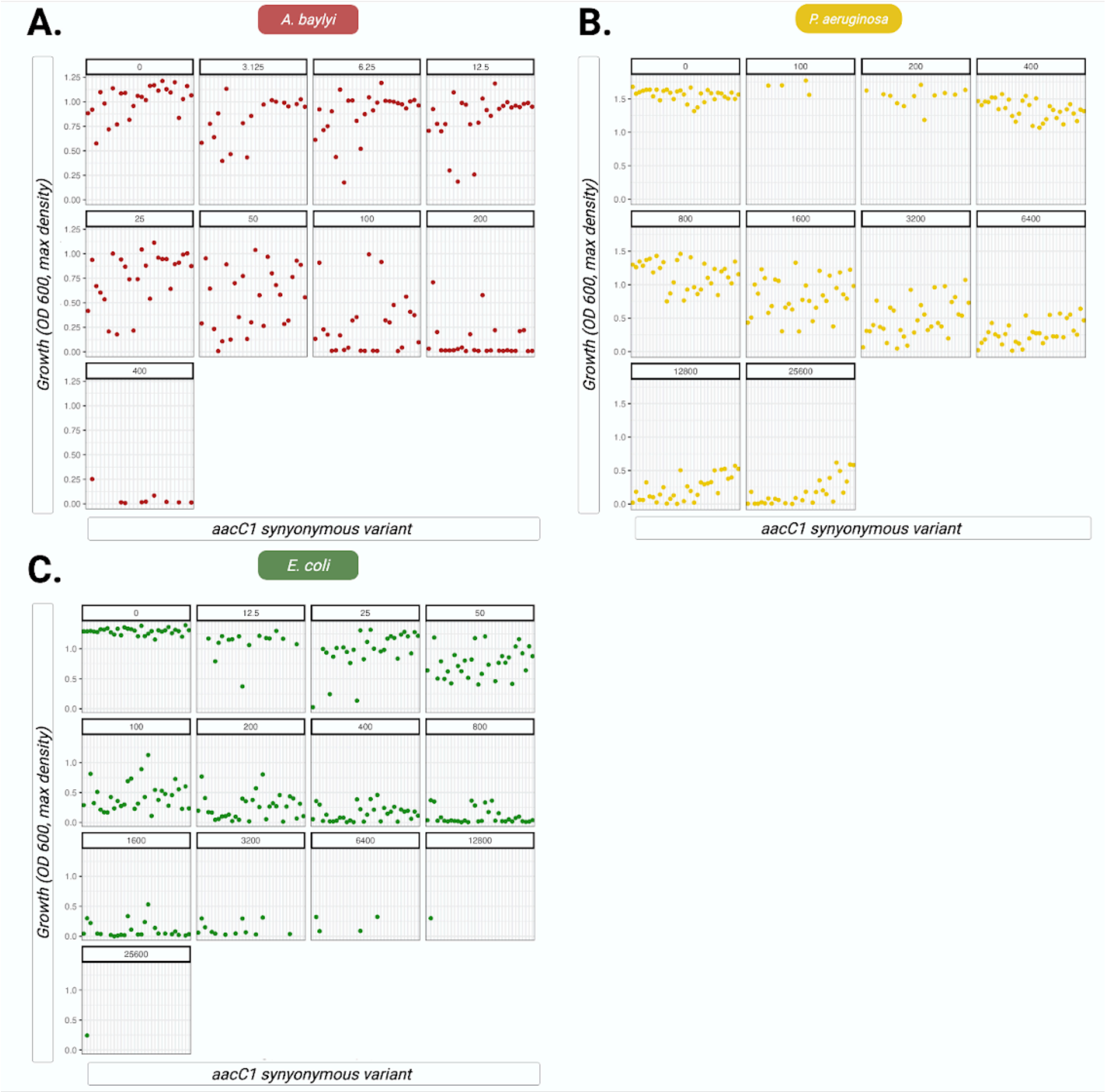


**Figure S5. Max OD 600.** Carrying capacity calculated as OD 600 max density for each variant in each gentamicin concentration. Variants are ordered from 1-32 on the x avis. The wt variant is V19. **A.** *A. baylyi.* Note that the graphs do not contain V3, V4, V12, V14, V20 or V32. **B.** *E. coli* **C.** *P. aeruginosa.* Max OD 600 individual sample data is available in supplementary file S24-S26.

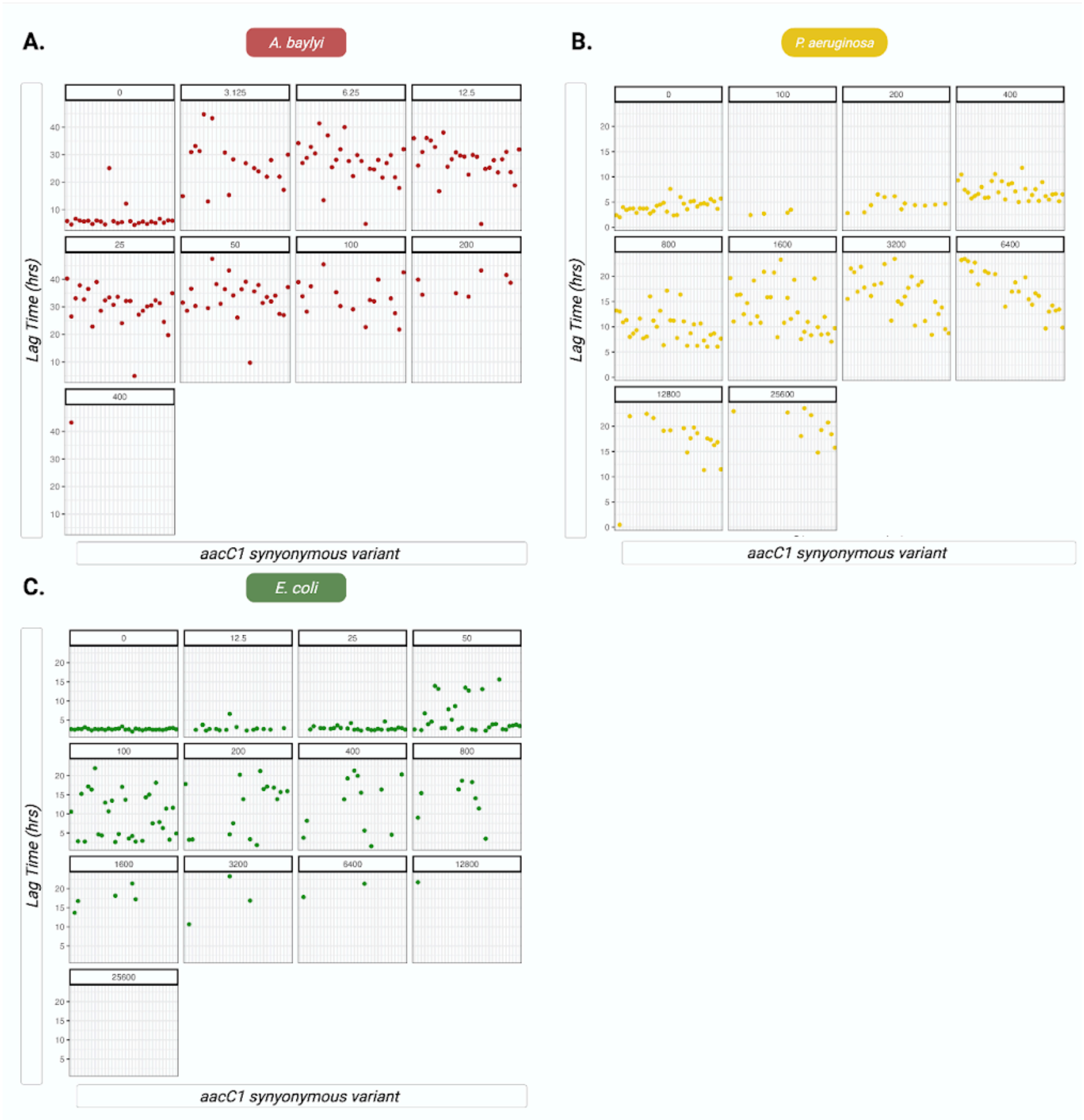


**Figure S6. Lag Time** Lag time calculated as the time needed for the bacterial culture to reach an OD 600 of 0.3. Variants are ordered from 1-32 on the x avis. The wt variant is V19. **A.** *A. baylyi*. Note that the graphs do not contain V3, V4, V12, V14, V20 or V32. *A. baylyi* cultures were grown for 48hrs. **B.** *E. coli*. **C.** *P. aeruginosa*. Max OD 600 individual sample data is available in supplementary file S24-S26.

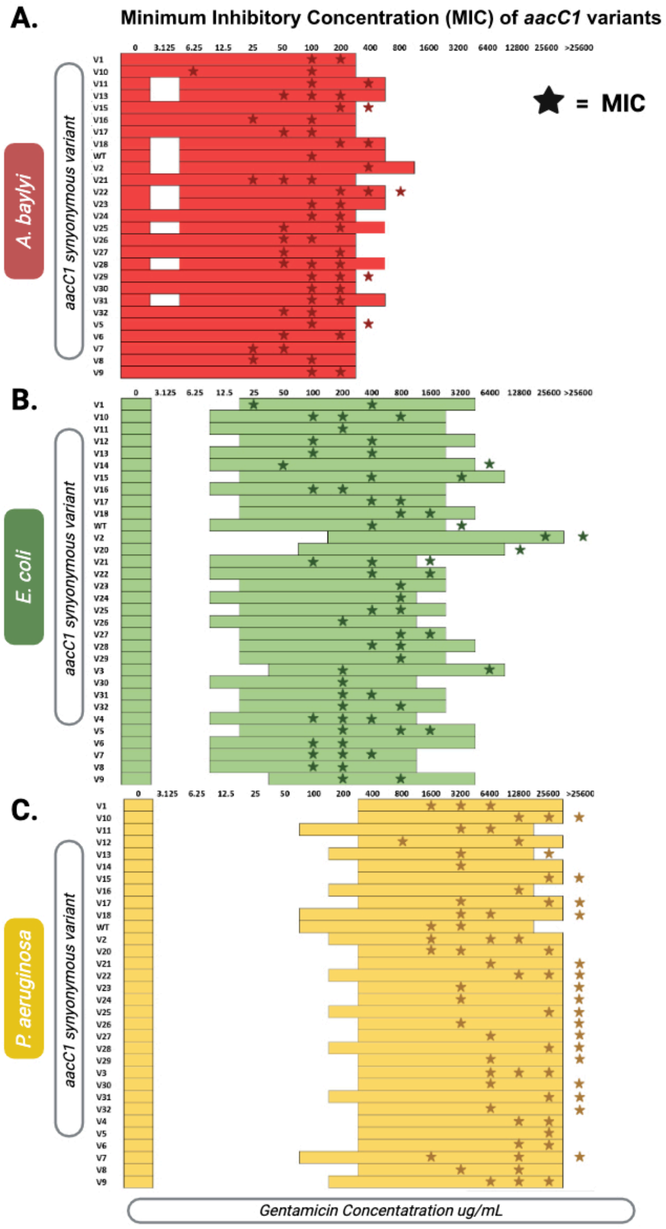


**Figure S7: Minimum Inhibitory Concentrations of *aacC1* variants:** Two-fold increases in gentamicin concentration are displayed on the x axis. aacC1 variants are represented on the y axis. Gentamicin concentration in which growth curves were performed are colored according to species. Non-measured concentrations are left blank. The MIC for each growth curve replicate is represented by a coloured star. MIC is defined here as the first concentration at which there is no visible growth (<0.1 OD 600). MIC stars outside of the coloured boxes represent samples that grew (>0.1 OD 600) in the respective highest measured gentamicin concentration. The next highest concentration is taken as the MIC and should be read as >Highest Measured Concentration (ex. *A. baylyi* V15 MIC = >200µg/mL gentamicin). ***A.*** *A. baylyi.* ***B.*** *E. coli.* **C .** *P. aeruginosa.*

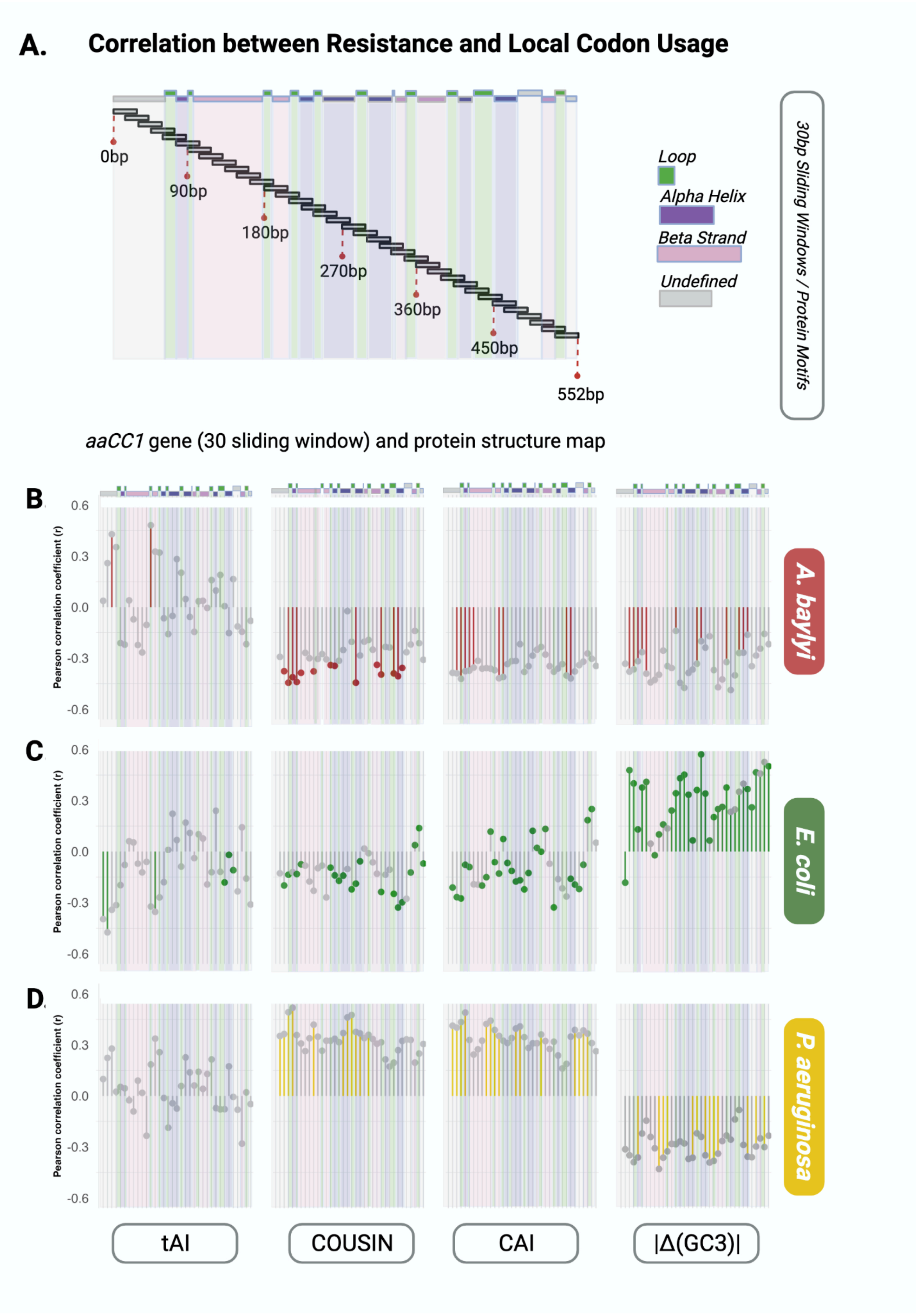

A.
Correlation between Resistance and Local Codon Usage
0bp
90bp
180bp
270bp
360bp
450bp
552bp
Loop
Alpha Helix
Beta Strand
Undefined
30bp Sliding Windows / Protein Motifs
aaCC1 gene (30 sliding window) and protein structure map
B.
C.
D.
Pearson correlation coefficient (r)
0.6
0.3
0.0
-0.3
-0.6
A. baylyi
E. coli
P. aeruginosa
tAI
COUSIN
CAI
|Δ(GC3)|

**Figure S8: Local codon usage and resistance (including Benjamini-Hochberg False Discovery Rate (FDR) corrections A.** Pearson correlation between resistance (IC50 AUC) and tAI at 30bp sized sliding windows (sliding every 15bps) of the *aacC1* gene, represented by r on the y axis. Lollypop graph icons represent the r value for each of the 35 sliding windows, ordered from 0-30 bp to 525-552 bp. Window 0-30bp is common to all *aacC1* variant sequences and is thus excluded from correlation tests and left blank. Non-significant correlations (p-value <0.05) are represented in grey. Samples with a p-value <0.05 are represented by a coloured lollipop stalk. Samples remaining after BH FDR are represented by a coloured lollipop head. **B.** Pearson correlation between resistance (IC50 AUC) and COUSIN **C.** Pearson correlation between resistance (IC50 AUC) and CAI. **D.** Pearson correlation between resistance (IC50 AUC) and |Δ(GC3)|. **E.** Protein structural elements, i.e. alpha helices, beta sheets and loops are represented in purple, pink and green, respectively. The positions of these elements within the coding sequence are displayed above the sliding window. Undefined elements are represented in grey.

| Codon | AminoAcid | *P.aeruginosa* | *A.baylyi* | *E.coli* | *B.cereus* |
|---|---|---|---|---|---|
| TTT | F | 0.10 | 1.61 | 1.15 | 1.41 |
| TTC | F | 1.90 | 0.39 | 0.85 | 0.59 |
| TTA | L | 0.02 | 1.78 | 0.79 | 3.10 |
| TTG | L | 0.43 | 1.26 | 0.78 | 0.62 |
| CTT | L | 0.15 | 1.02 | 0.62 | 1.15 |
| CTC | L | 1.34 | 0.49 | 0.62 | 0.26 |
| CTA | L | 0.07 | 0.50 | 0.22 | 0.67 |
| CTG | L | 3.99 | 0.96 | 2.96 | 0.20 |
| ATT | I | 0.21 | 2.04 | 1.52 | 1.87 |
| ATC | I | 2.72 | 0.70 | 1.26 | 0.48 |
| ATA | I | 0.07 | 0.26 | 0.22 | 0.65 |
| ATG | M | 1.00 | 1.00 | 1.00 | 1.00 |
| GTT | V | 0.16 | 1.23 | 1.04 | 1.42 |
| GTC | V | 1.67 | 0.74 | 0.86 | 0.31 |
| GTA | V | 0.23 | 0.99 | 0.62 | 1.69 |
| GTG | V | 1.93 | 1.04 | 1.48 | 0.59 |
| AGT | S | 0.29 | 1.47 | 0.91 | 1.50 |
| AGC | S | 2.80 | 0.82 | 1.65 | 0.60 |
| TCT | S | 0.10 | 1.35 | 0.88 | 1.58 |
| TCC | S | 1.31 | 0.33 | 0.89 | 0.32 |
| TCA | S | 0.07 | 1.39 | 0.75 | 1.53 |
| TCG | S | 1.42 | 0.63 | 0.92 | 0.47 |
| CCT | P | 0.18 | 1.37 | 0.64 | 1.09 |
| CCC | P | 1.03 | 0.40 | 0.50 | 0.14 |
| CCA | P | 0.18 | 1.69 | 0.77 | 1.93 |
| CCG | P | 2.61 | 0.53 | 2.09 | 0.83 |
| ACT | T | 0.17 | 1.03 | 0.67 | 0.92 |

**Table S1: Relative Synonymous Codon Usage (RSCU) of each codon in each of the four species.** RSCU is calculated as the frequency of use of a synonymous codon in relation to all of the codons encoding the same amino acid. The sum of RSCU values for each amino acid equals the number of synonymous codons for that amino acid.
